# The hidden costs of dietary restriction: implications for its evolutionary and mechanistic origins

**DOI:** 10.1101/533711

**Authors:** Andrew W McCracken, Gracie Adams, Laura Hartshorne, Marc Tatar, Mirre J. P. Simons

## Abstract

Dietary restriction (DR) consistently and universally extends health- and lifespan across taxa. Despite considerable research, precise and universal mechanisms of DR have not been identified, limiting the translational potential of its beneficial outcomes to humans. In biomedical science, DR’s effects are interpreted as stimulating pro-longevity molecular pathways. This rationale is guided by the conviction that DR evolved as an adaptive, pro-longevity physiological response to restricted food supply. Current evolutionary theory states that organisms should invest in their soma more heavily during periods of DR, and, when their resource availability improves, should outcompete age-matched rich-fed controls in survival and/or reproduction. Here we present a formal test of this key prediction utilising a large-scale demographic approach detailing mortality and fecundity in Drosophila melanogaster fed alternating dietary regimes (N > 66,000 flies across 11 genetic lines). Our experiments reveal substantial, unexpected mortality costs when returning to a rich diet following periods of DR, in direct contrast to the predictions from current evolutionary theory of DR. The physiological effects of DR should therefore not be interpreted as being intrinsically pro-longevity, acting through increased investment in somatic maintenance. We suggest DR’s effects could alternatively be considered an escape from costs incurred under nutrient-rich conditions, in addition to novel, discrete costs associated with DR. Our results therefore question the relevance of DR’s current evolutionary explanation in interpreting its mechanistic basis.

## Introduction

Ageing has attracted extensive scientific interest, both from a fundamental and biomedical perspective. Dietary restriction (DR) universally extends health- and lifespan across taxa, from baker’s yeast to mice (Fontana and Partridge, 2015). The reduction of total calories - or restriction of macronutrients, such as protein - extends lifespan reliably (Min *et al.*, 2007; Lee *et al.*, 2008; Solon-Biet *et al.*, 2014; Jensen *et al.*, 2015). Although the precise universal mechanisms that connect diet to ageing remain elusive, translation of DR’s health benefits to human medicine is deemed possible (Dirks and Leeuwenburgh, 2006; Balasubramanian, Howell and Anderson, 2017). The widespread assumption of DR’s translational potential originates from the notion that DR’s beneficial effects are facilitated by shared evolutionary conserved mechanisms, as beneficial effects of DR are observed across taxa. Indeed, experiments on our close evolutionary relatives, rhesus monkeys (*Macaca mulatta*) have demonstrated that DR could be translational (Mattison *et al.*, 2017). Still, the mechanisms by which these benefits are accrued physiologically may be wholly disparate, as no single genetic or pharmaceutical manipulation mimicking the benefits of DR across model organisms exists (Selman, 2014). Mechanistic insight will be key, since DR as a lifestyle intervention has limited scope, given the degree of self-restraint required. It is therefore warranted to direct scrutiny towards the evolutionary theory of DR, since it underpins the assumed universality of physiological mechanisms by which DR confers health benefits.

Shared universal mechanisms can only be inferred from the ubiquity of the DR longevity response, when the selection pressures responsible for such evolutionary conservation are understood. The DR response itself may have evolved once, and mechanisms might be conserved. Alternatively, DR could have undergone convergent evolution, either using similar mechanisms - or by adopting alternative ones (Mair and Dillin, 2008). These evolutionary scenarios provide distinct predictions as to how informative mechanistic research in other animals will prove for human medicine. Current evolutionary theory on DR is limited, and its elemental phenotypic predictions have undergone minimal empirical examination (Zajitschek *et al.*, 2018). The DR effect itself is interpreted as an evolved, adaptive, pro-longevity physiological response to limiting food availability (Holliday, 1989). Life-history theory - a central tenet of evolutionary biology - states resources are limited, and thus predicts trade-offs between reproduction (Höglund, Sheldon and Hoglund, 1998) and survival (Stearns, 1989), even in nutrient-rich environments. As such, DR presents an enigma: why do organisms live longer on a constrained energy budget?

The currently accepted evolutionary model for DR (Shanley and Kirkwood, 2000) uses a life-history perspective on ageing - the Disposable Soma Theory (DS) that predicts a trade-off between investment into reproduction and somatic maintenance (Kirkwood, 1977; Kowald and Kirkwood, 2016) - to explain this enigma. The model proposes that below a certain resource threshold, organisms will reallocate energy almost exclusively towards somatic maintenance (Fig.1). In certain ecological situations (e.g. severely reduced juvenile survival, or when the energy budget is lower than the initial costs (Jönsson, 1997), or the cost of one unit, of reproduction) investment into reproduction will cease to yield fitness. The optimal, fitness-maximising strategy under these harsh conditions would be to terminate investment into reproduction and utilise this energy to gain fitness when conditions improve. Crucially, this life-history strategy could favour an increase in energy devoted to maintenance and repair - allowing organisms to survive periodic bouts of famine with an intact (or superior) soma (Shanley and Kirkwood, 2000; Kirkwood and Shanley, 2005). This ‘somatic maintenance response’ has been presumed to be the primary causative agent in the pro-longevity DR response (Kirkwood and Austad, 2000; Masoro, 2000; Ingram *et al.*, 2006; Speakman and Mitchell, 2011; Fontana and Partridge, 2015).

**Fig 1.**
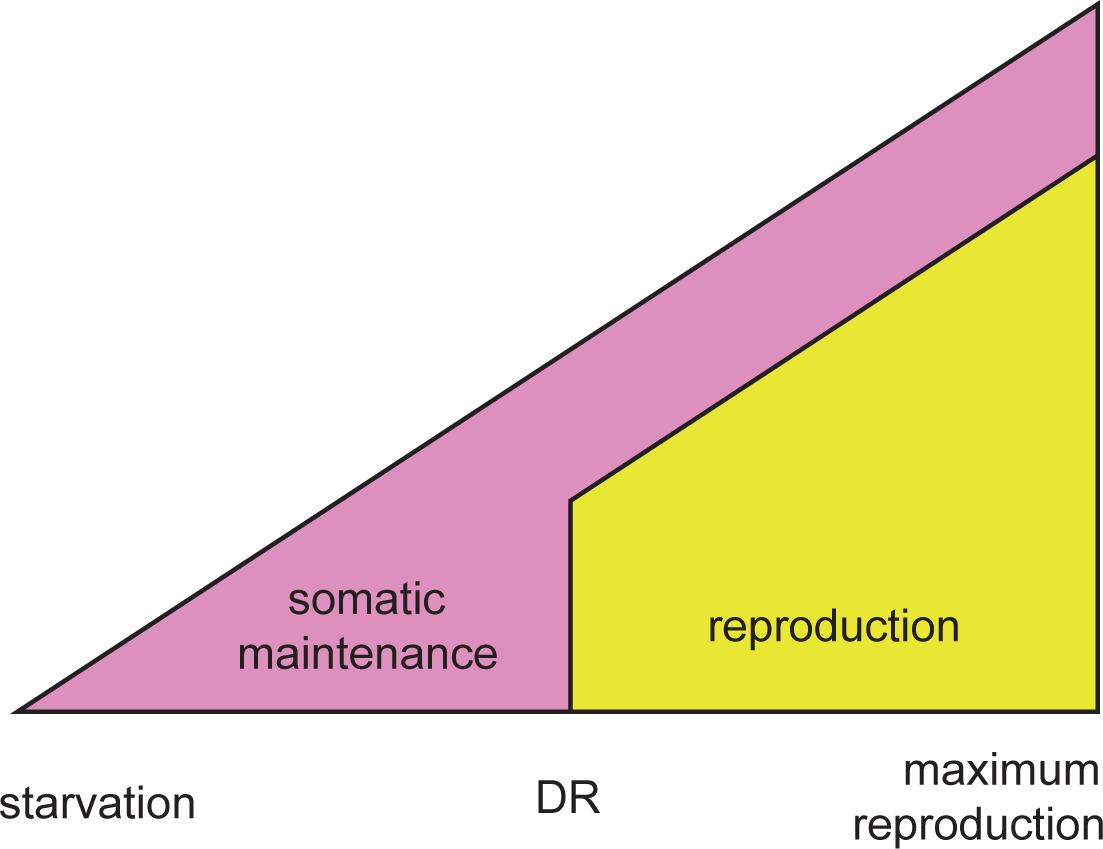
Schematic of the evolutionary model of DR. Resource availability is varied from left to right, from very low (where starvation would occur) to very high (where maximum reproduction would occur). The theoretical optimal allocation to somatic maintenance (pink) versus reproduction (yellow) is depicted at a given resource availability. When resource availability decreases, investment in both somatic maintenance and reproduction is reduced, until a threshold is met. Below this point resources are so scarce that investment in reproduction does not yield a fitness return. This could occur when offspring produced cannot recruit into the population due to the harsh resource environment, or because the capital (start-up) costs of breeding cannot be met. Here, investment in reproduction is lost and is wholly allocated to somatic maintenance. It is this evolved resource allocation decision to invest into somatic maintenance under DR conditions, that is thought to underlie lifespan extension under DR.

This attractive evolutionary rationale has given credibility to the assumption that physiological changes in the DR animal are inherently pro-longevity. For example, transcriptomic upregulation of, what could be, maintenance processes under DR, have lent credence to this hypothesis (Lee *et al.*, 1999; Kirkwood and Austad, 2000; Pletcher, Libert and Skorupa, 2005; Whitaker *et al.*, 2014). Directionality is often ambiguous, however; downregulation of DNA repair under DR could be interpreted as either a reduction in DNA damage generation, or reduced investment into repair (Lee *et al.*, 1999; Pletcher, Libert and Skorupa, 2005). In other words, a simpler rationale is often neglected: the surge of ‘maintenance and repair’ gene expression could be a ‘mere’ stress response to metabolic disruption. The health benefits observed under DR might originate from a passive response - one not necessarily evolved as an adaptive regulatory response to diet. Under these circumstances, lifespan extension could be a ‘simple’ correlated response to currently unknown, but strongly conserved, physiology. For example, the limitation of metabolic rate or reduction in specific metabolites as a direct consequence of DR could reduce conserved associated physiological dysfunction, and thereby extend lifespan. The negative physiological effects dietary restricted organisms suffer, e.g. compromised immune function (Kristan, 2008) and cold intolerance (Adler and Bonduriansky, 2014), could arise from a similar passive response, and are not necessarily the result of a regulated trade-off. DR is sometimes considered a hormetic response - mild stress, resulting in the stimulation of conserved cellular reactions leading to beneficial health (Rattan, 2008) – which would be a similar example of a passive response. One example of such a response is the activation of heat shock proteins, which show only very transient expression (Li and Duncan, 2008), but long-lasting effects on life expectancy (Tatar, Khazaeli and Curtsinger, 1997).

The distinction between more passive correlated responses versus adaptive programmed pro-longevity responses will be key to identifying the basic mechanisms of DR and develop translation to humans. The widely-accepted evolutionary model of DR supports an adaptive phenotypic response and provides a key prediction: organisms should invest in their soma more heavily during periods of DR, and, when their resource availability improves, should outcompete age-matched rich-fed controls in survival and/or reproduction. Here we provide an experimental test of this prediction, utilising a large-scale demographic approach detailing mortality and fecundity in *Drosophila* melanogaster fed different dietary regimes. Our results revealed substantial mortality and fecundity costs when returning to a rich diet after a period of DR, falsifying the key prediction provided by the evolutionary biology of DR. These effects were independent of genotype, duration of DR, number of dietary fluctuations, access to water, and microbiota abundance. Our results therefore suggest that the effects of DR are not intrinsically pro-longevity, and could be considered an escape from costs incurred under nutrient-rich conditions in addition to novel, discrete costs associated with restricting dietary protein. These insights question the relevance of current evolutionary explanations of DR in guiding biomedical research into its mechanisms. Our alternative paradigm - a passive, not necessarily directly adaptive response to DR - gives renewed credibility to a range of mechanistic hypotheses of DR: hormesis (Masoro, 2005; Sinclair, 2005), a reduction in metabolism causing reduced oxidative damage generation (Masoro, 2005; Mair and Dillin, 2008; Redman *et al.*, 2018) and improved mitochondrial functioning (Weir *et al.*, 2017), or a reduction of waste products from specific metabolic pathways (Hipkiss, 2006).

## Results

### Hidden costs of Dietary Restriction

Our first set of diet experiments were conducted using the wild-type inbred lineage DGRP-195, comprising 11,084 individual deaths (Table S1). DR imposed continuously throughout adult life resulted in a significant reduction in mortality rate (Fig. 2; Table S1,2; P < 0.001, 3 times lower hazard). In addition, switching flies to DR at older ages instantly reduced mortality levels to the levels of flies that had experienced continuous DR (‘short reverse-switch’; Fig. 2D; Table S1). Such mortality amnesia - a complete absence of historic diet effects - has been reported previously in flies (Good and Tatar, 2001; Mair *et al.*, 2003).

Our expectation, based on the current evolutionary model of DR, was that if flies were returned to rich food conditions after a period of DR, they would have a superior soma compared to flies that experienced rich food continuously. Energy reallocated from reproduction towards somatic maintenance should result in higher fitness and enhanced longevity (Fig. 1). In contrast, our ‘long-switch’ treatment resulted in a substantial increase to mortality risk compared to flies kept on a rich diet throughout life (Fig. 2A; Table S1; P < 0.001, 3.7 times higher hazard). Mortality peaked immediately (within 48h; 5.1 times higher hazard) after the switch from a restricted to a rich diet. The magnitude of this mortality difference decreased slowly thereafter, resulting in no difference between the continuous rich diet and the long-switch treatments after eight days (Fig. 2A; Table S3; P < 0.001).

**Fig 2.**
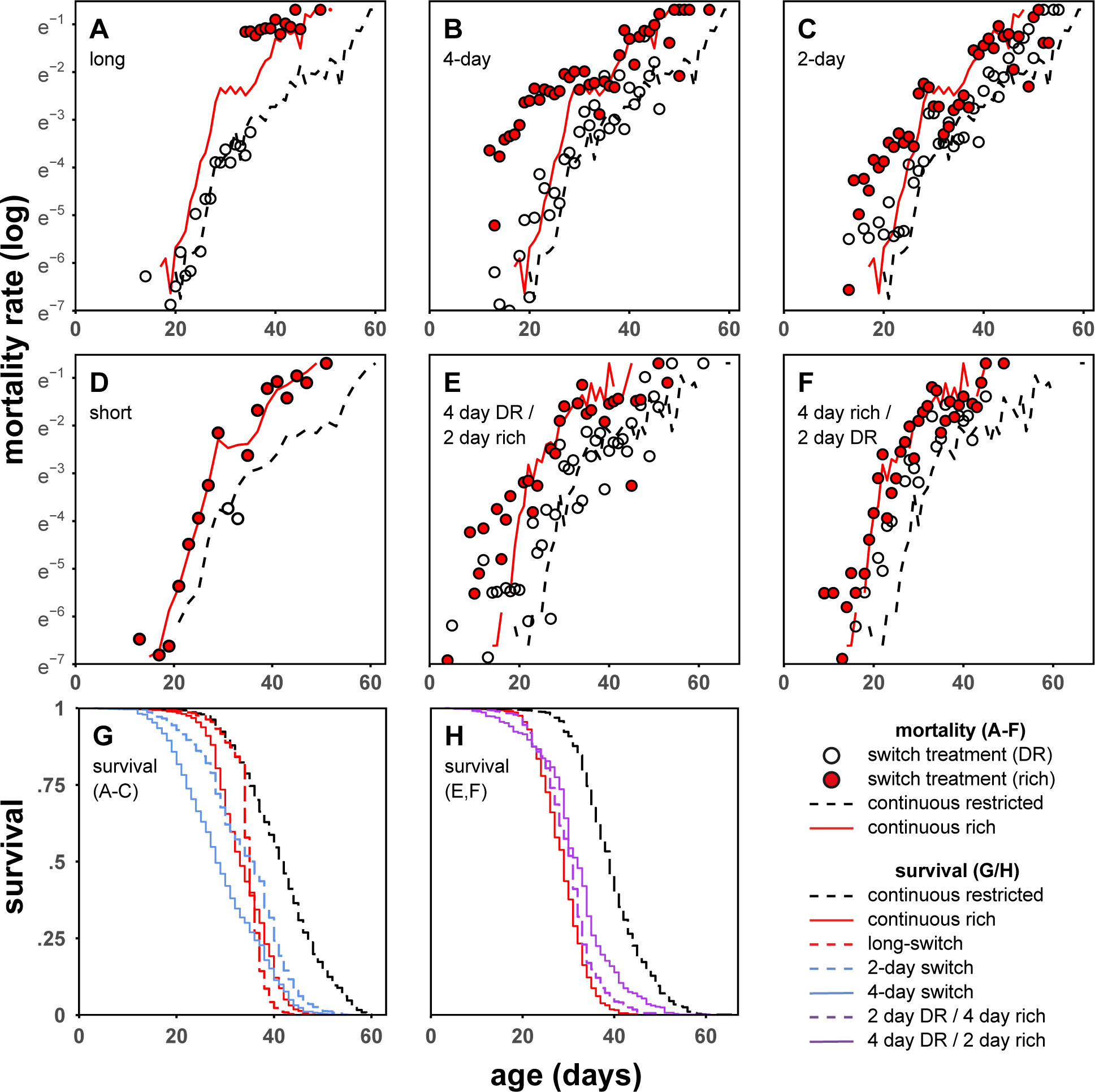
The effect of different dietary regimes on age-specific mortality risk in DGRP-195. N = 19,086 total. N = 995-3,769 per treatment. Mortality risk at continuous rich and restricted diets are plotted as lines. Dietary switch treatments are plotted as points. The exacerbation of mortality due to switch phenotypes is observable as the difference between mortality at continuous rich diet (red line), and mortality of switch treatment when on a rich diet (red points). A – long-switch. When returning to a rich diet after a long period of DR, mortality is exacerbated compared to flies fed a rich diet continuously. B – 4-day switch. Switching from a DR to a rich diet repeatedly every four days, increases mortality on rich diets compared to continuously rich fed flies. Flies are still able to modulate their mortality in response to DR, even when diet fluctuates rapidly. C – 2-day switch. Mortality on rich diets is only mildly increased and flies still respond to DR even when it is only imposed for two days. D – short reverse-switch. After a long period on a rich diet, DR for 4 days returns flies to mortality of continuous DR. The x-axis of panel D is age-adjusted to correct for age differences (1-3 days) at the time of the diet switch for illustration purposes only. E – 4-day DR, 2-day rich switch (4-to-2 day switch). Flies respond to DR, but encounter a slightly blunted effect compared to continuous DR. F – 4-day rich, 2-day DR switch (4-to-2 day switch). The effect of DR is reduced when imposed for 2 days following 4 days on a rich diet. G – survival plot of panels A-C with associated continuous diet controls. Total survival of both the 4-day switching dietary regime and the long-switch is lowered compared to continuously rich diets, despite flies spending a considerable extent of their lives on restricted diets. Flies on DR outlive all other categories. H – survival plot of panels E/F with associated continuous diet controls. Despite spending up to two-thirds of their lives on DR in these asymmetrical regimes, survival benefits are modest, compared to continuous DR. Dietary switch treatments contain daily time-points (dots) for the dietary switch treatments, as treatments were mirrored and balanced, with half of flies starting on DR, and half on rich diets.

### Repeated diet switching

The long-switch dietary treatment may be dependent on several specific aspects of the imposed dietary regime, not necessarily falsifying the somatic maintenance hypothesis of DR. First, the effects of the long-switch treatment could be contingent upon the duration of prior exposure to DR. Indeed, it has been suggested that DR evolved as a response to relatively short, intermittent bouts of famine (Shanley and Kirkwood, 2000). Second, it has also been suggested that the longevity effect of DR itself may be selected for at relatively young ages (Shanley and Kirkwood, 2000). Thus, it was possible that young flies would not show the heightened mortality we observed. Third, it could be that sudden changes in diet *per se* is harmful. To test these three potential confounds we used short recurring bouts of DR, alternating between a rich and a DR diet every four days (‘4-day switch’). In this dietary regime, mortality on the rich diet compared to the continuous rich diet was similarly exacerbated (Fig. 2B; Table S1, 2; P < 0.001, 2.4 times higher hazard). This 4-day switch dietary regime also allowed us to examine whether flies were able to instantly and repeatedly modulate their mortality risk in response to diet, similar to the short reverse-switch treatment (Fig. 2D). Flies indeed modulated their mortality in response to the diet they were currently fed, with a degree of surprising immediacy. Mortality risk on DR, within the 4-day switch regime, repeatedly decreased to levels similar to that of flies continuously exposed to a restricted diet (Fig. 2B; Table S1). Nonetheless, mortality risk during these periods of DR imposition was significantly higher than that of continuous DR-treated flies (Table S1; P < 0.001, 1.6 times higher hazard). We suggest this increase in mortality seen on DR in the 4-day switch treatment is due to either accrued physiological costs or more probable, a carry-over of deaths directly resulting from the rich diet, but recorded on the DR diet.

### Mortality costs depend on the duration of Dietary Restriction

A closer examination of the timing of mortality within the 4-day switching paradigm showed that the mortality response was strongest in the second 48 hrs after exposure to both DR and rich diets (Table S4; P < 0.001). This suggests a period of acclimation to both DR and rich diets is necessary before their physiological effects are fully realised. To test the importance of the duration of exposure to DR and rich diets for the mortality phenotypes observed, further dietary regimes were used. First, switching from DR to rich conditions was carried out at increased frequency - alternating every 2 days (‘2-day switch’; Table S1, 2). This 2-day switch dietary regime confirmed that sustained exposure to the diets (longer than 2 days) was required to cause the mortality phenotypes observed. On a rich diet, the 2-day switch regime showed slightly higher mortality compared to the continuous rich diet (Fig. 2C; hazard = 1.1, P < 0.05) and mortality on DR in the 2-day switch regime did not reduce to the levels seen in continuously dietary restricted flies (Fig. 2C; hazard = 1.3, P < 0.001). Together these diet-specific mortality effects resulted in an overall lifespan extension in the 2-day switch regime (Fig. 2G; Table S2; P < 0.001). As flies spend an equal amount of time on DR or rich diets in the 2-day switch regime: the reduction of mortality under DR can be considered to be relatively more rapid than the induction of exacerbated mortality on rich food (after a period of DR). We reasoned that the exacerbation of mortality on rich food either requires an extended period on restricted or rich food. To test this directly, asymmetrical dietary regimes were used.

In this additional set of experiments, we combined the 4-day and 2-day switching regimes: treatments were comprised of 4 days on either a DR or rich diet, followed by 2 days on the other (‘4-to-2 day switch’). Similar to the 4-day switch, this dietary regime was repeated sequentially. These ‘4-to-2’ regimes showed no marked increase in mortality on the rich diet compared to flies on a continuous rich diet (Fig. 2E, F; Table S5). Relative to a continuous DR treatment, the effect of DR within this paradigm was markedly reduced, especially when flies were restricted for 2 days only (Fig. 2F; Table S5). This reduction in the mortality response to DR in the ‘4-to-2’ regimes amounted to a marked reduction in the total longevity extension achieved when compared to continuous DR. When flies spend two-thirds of their lives on DR lifespan was only extended by half (compared to continuous DR), and only a quarter when flies spend one-third of their lives on DR (Fig.2E, F; Table S6). These experiments thus again suggest a period exceeding 2 days on either diet is required to induce marked mortality effects.

Note that within the long-switch treatment, the mortality exacerbation observable on rich food was strongest within the first 2-day interval (Fig 2A; Table S3). Additionally, our short reverse-switch induced a full DR response - mortality amnesia - within 2 days (Fig. 2D; Table S1). Moreover, the ameliorated mortality exacerbation of our additional switch experiments (2-day, and 4-to-2 day switches) strongly suggest that the sudden dietary perturbations themselves are not the cause of premature mortality in our switching regimes. From these combined results we therefore tentatively conclude that the additional mortality costs observable on a rich diet are contingent upon the prior duration of DR.

### Genetic variance

To eliminate the possibility that the dietary responses described above (in DGRP-195) were the result of rare genetic effects, we performed the same dietary perturbations in a panel of randomly selected genotypes (DGRP-105; 136; 195; 217; 239; 335; 362; 441; 705; 707; 853). Across our panel, we detected an increase in longevity under DR conditions (Fig. 3, 4; additive model, DR hazard = −0.21 ± 0.08, P < 0.001). There were considerable genetic effects in response to diet however (interaction model: X^2^ = 204.8 (df=10), P < 0.001), with some genotypes showing elevated mortality under restricted-diet conditions, compared to continuously-fed rich diet flies (Fig. 3, 4). This degree of variation in response to DR can be explained by genetic variation in the reaction norm to diet. A particular combination of a restricted and rich diet will not always induce the same longevity response in a range of genotypes (Lee *et al.*, 2008; Tatar, 2011; Jensen *et al.*, 2015).

**Fig 3.**
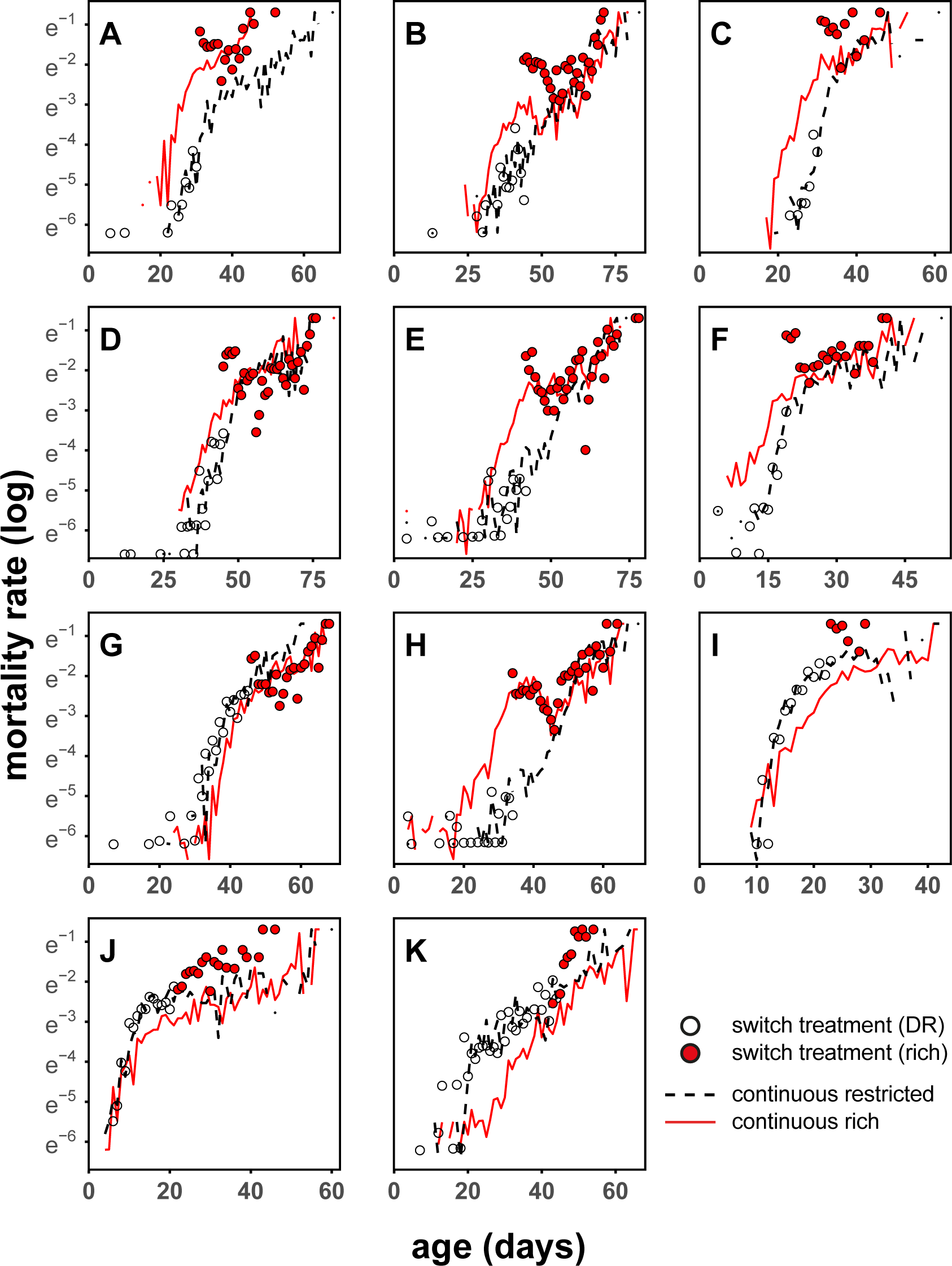
Long-switch treatment in a panel of 11 DGRP genotypes. A – 195; B – 105; C – 217; D – 441; E – 705; F – 707; G – 136; H – 362; I – 239; J – 335; K – 853. N = 29,702 total; ∼2,725 per genotype; 13,375 for continuous rich treatments, and ∼8,170 each for the two other treatments. The dietary switch for the long-switch treatment group occurred at 45-65% of continuous rich treatment flies. All panels contain daily time-points as in Fig.2. Exposure to a high nutrient diet after a period of DR resulted in marked increase in mortality compared to a continuous rich diet in all lines (9 out of 11 significant). There was genetic variation in this response with DGRP-136 (G) and DGRP-362 (H) showing the smallest effects. This marked overshoot was not contingent upon DR extending lifespan. Lines that showed ‘starvation’ on a DR diet still showed significant overshoots when they were switched to a rich diet, where recovery from starvation was expected, even when compared to continuous DR diets (I, J, K).

**Fig 4.**
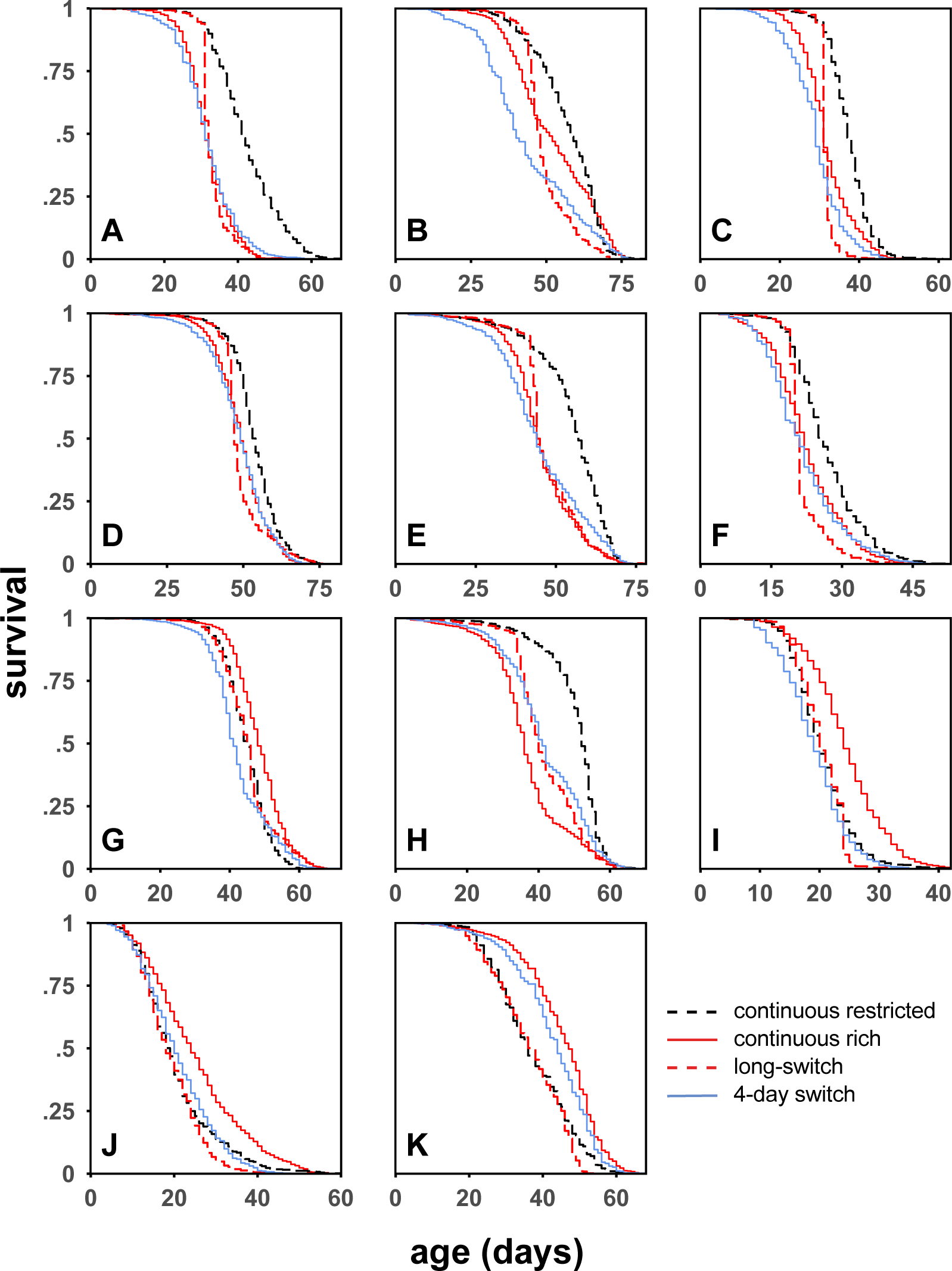
Survival curves of DGRP panel for both dietary regimes. A – 195; B – 105; C – 217; D – 441; E – 705; F – 707; G – 136; H – 362; I – 239; J – 335; K – 853. Total survival on the different dietary regimes across the genetic panel tested. Rich diets after a period of DR resulted in such an increase in mortality, that total survival of the cohort was lower (or equal to) those fed a continuous rich diet for their whole life (A-F). N = 37,897 total; ∼3,450 per genotype; 13,375 for continuous rich treatments, and ∼8,170 for all other treatments.

We tested genetic variance in the response to the long-switch treatment using age-dependent interval based models. Across genotypes, exposure to the rich diet after a period of DR (long-switch) resulted in exacerbated mortality, exceeding that of flies fed a rich diet for their whole lives (Fig. 3; additive model, hazard = 0.997 ± 0.056, P < 0.001). There was significant genetic variance for this trait (χ^2^ = 124 (df=10), P < 0.001). Still, all genotypes showed a sudden and marked mortality overshoot, compared to a continuous rich diet, following a switch from DR to high nutrient conditions (Fig. 3; Table S7, 8; 9 of 11 significant; range: 1.12 to 5.21 times hazard).

Alternating the diet from DR to rich every 4 days decreased longevity compared to the continuous rich diet, across the genetic panel (additive non-interval based model, hazard = 0.24 ± 0.047, P < 0.001). Again, we found significant genetic variance for the response to this dietary regime (χ^2^ = 117 (df=10), P < 0.001). Lines differed in their responses: 5 of the 11 showed marked decreases in survival; 1 showed an increase in survival, and the remaining 5 showed statistically non-significant effects (Table S8). Interval-based models showed that mortality rates increased at the rich diets following a period of DR, as in the long-switch, in all lines (significant in 7 of 11; Table S10,11). There was a modest positive genetic correlation in the increase in mortality induced by the long-switch and 4-day-switch dietary regimes (correlation of coefficients from Table S9, 11; r_s_ = 0.45, P = 0.17), suggesting these dietary phenotypes originate from similar physiology.

### Hidden costs: independent of a pro-longevity DR response

Our restricted diet unexpectedly induced a putative starvation response - observable as an increased mortality rate - in four lines (136, 239, 335, 853; Fig. 3, 4; Table S7, 9, 10, 11). These contrasting responses to DR serendipitously allowed us to see whether the dietary switching phenotypes were contingent on the direction of the DR response. Surprisingly, when lines that showed starvation were refed on a rich diet (long-switch), mortality did not decrease, but increased (Table S7, 8; 3 out of 4 showed a significant increase), even beyond the heightened mortality seen on DR (Fig.3, Table S12).Similarly, within the 4-day switching regime, mortality risk was exacerbated at a rich diet. The pattern of mortality even reversed, compared to individuals fed diets continuously, with lines now showing a putative DR-longevity response within the 4-day switch dietary regime (Fig. S1; Table S10, 11, 13). These outcomes were particularly remarkable since exposure to a richer diet was expected to rescue the starvation response. In essence however, upon a return to the diet where recovery would be expected, individual mortality risk surged even higher.

### Cost of mortality not compensated for by fecundity increase

We recognised our results would not necessarily discredit the evolutionary model of DR should the observed costs in mortality be compensated fully, or partially, by an increase in fecundity. Egg production across the DGRP panel experiment was measured from vials in each dietary regime and expressed both as a total count (age-specific fitness of the population; Fig. S2, 3; Table S14, 16) or eggs per fly (age-specific reproductive output, corrected for mortality differences; Fig. S2, 3; Table S15, 17). All lines responded strongly to DR, by reducing reproductive output. Within the 4-day switching paradigm, the restricted diet also induced a rapid reduction in fecundity (Fig. S3; Table S16, 17). As with the mortality response, genetic lines responded somewhat differently in fecundity in response to the dietary treatments (long-switch: F=57 (df=2), P < 0.001; 4-day switch: χ^2^ = 187 (df=9), P < 0.001). However, in both metrics, our switching diets underperformed in comparison to the continuous rich diet (Fig. S2, 3; Table S14-17), confirming our mortality phenotypes were not compensated by higher fecundity upon a return to nutrient rich conditions.

### Mortality phenotypes were not contingent on condition of the microbiome, social housing, water or sex

A switch to rich diets after a sustained period of DR (long-switch) still resulted in an increase of mortality when flies were treated with antibiotics (Table S18; P < 0.001), provided additional water (Table S19; P = 0.002), or when mortality was assessed in isolation (Table S20; P = 0.014). Males responded, similar to females, by increasing mortality on rich diets if this was preceded by 4 days of DR (4 day switch, Table S21; P = 0.001, long-switch not tested).

## Discussion

DR has been tested across multiple species and the resulting lifespan extension has consistently - with very few exceptions (Adler and Bonduriansky, 2014) - been interpreted as provoking anti-ageing, pro-longevity physiology. This interpretation is based on the widely-accepted evolutionary theory of DR (Shanley and Kirkwood, 2000; Kirkwood and Shanley, 2005) which predicts that during periods of DR, investment in somatic maintenance is actively increased, to await better times when fitness can be gained. In contrast, we find that periods of DR did not result in a superior soma, and instead resulted in large increases in mortality, and reductions in fecundity, when nutrient availability returned to plentiful. Our results question the current explanation of DR’s evolutionary origins, and thereby their relevance in interpreting DR’s mechanistic origins.

Other studies have raised similar concerns but have only very rarely measured the consequences of the relevant life-history event: a period of DR followed by a period of rich food conditions. Direct measurement of investment into the soma using stable isotopes showed no increased investment under DR (O’Brien *et al.*, 2008). Experimental evolution across fifty generations under DR, failed to support the current evolutionary theory of DR (Zajitschek *et al.*, 2018). Further lack of support, we suggest, originates from the remarkably immediate reduction in mortality – a reduction in frailty, rather than actuarial ageing rate (Good and Tatar, 2001; Mair *et al.*, 2003; Simons, Koch and Verhulst, 2013) or historic physiological effects of diet (Selman and Hempenstall, 2012; Rusli *et al.*, 2018)– seen when flies are dietary restricted. A limited number of previous studies with *Drosophila* have shown such a response (Good and Tatar, 2001; Mair *et al.*, 2003). We confirmed these results (Fig. 2D), but also show for the first time that flies are capable of doing this repeatedly, in response to multiple switches in diet. Since DR does not slow ageing demographically, but results in an instant lowering of mortality - without any accrued beneficial effects - this is in itself evidence against increased somatic investment under DR (Simons, Koch and Verhulst, 2013; Garratt, Nakagawa and Simons, 2016).

In the reverse scenario, when flies resumed rich diets after DR, their performance was markedly lower than that of flies that were fed rich diets for their entire lives. Notably, this effect held, even when DR caused starvation (Tatar, 2011) - resulting in exacerbated mortality on the diet that should have provided an opportunity to refeed. Previous studies did not detect the same mortality costs in dietary regimes analogous to our long-switch (Mair *et al.*, 2003; Mair, Piper and Partridge, 2005), although in the raw non-smoothed data, some exacerbation of mortality can be seen in some conditions. There are a number of potential variables which could explain these differences. First, the duration of DR prior to a rich diet appears to be integral to inducing exacerbated mortality on rich diets (Fig. 2). Second, the existence and intensity of both the long-switch and 4-day switch phenotype, are genotype-dependent (Fig. 3, S1). This matter is further complicated by the lack of complete synchronicity between both phenotypes, across genotypes (Fig. 3, S1). Last, the longevity response to both a restricted diet, and the re-introduction of a rich one, may be contingent on the macronutrient composition of both (Lee *et al.*, 2008; Jensen *et al.*, 2015). Earlier work diluted media reducing both carbohydrates and protein (Mair *et al.*, 2003; Mair, Piper and Partridge, 2005), in contrast to our method of reducing yeast concentration, protein, alone.

Genotypes will differ in their longevity reaction norm to diet, rendering it impossible to know *a priori* whether a certain dietary composition constitutes the exact optimal longevity-directed diet (Tatar, 2011; Flatt, 2014). Genetic variation in the response to DR, reported in rodents (Liao *et al.*, 2010; Swindell, 2012; Mitchell *et al.*, 2016) and flies (Wilson *et al.*, 2017), might therefore not necessarily, or wholly, constitute variation in the physiological mechanisms that connect DR to ageing. We propose that our dietary phenotypes may also be contingent upon the direction and degree in which these diets deviate from the optimum, which may be one explanation for the dissimilarity of results observed in similar experiments. These considerations may also explain why the precise duration of DR is important, in line with the recent finding that the duration of starvation is critical in the lifespan extension generated via intermittent fasting (Catterson *et al.*, 2018). In addition, larval diet, timing and the order of how diets were fluctuated contributed to differential mortality observed when fluctuating diet (van den Heuvel *et al.*, 2014). Interestingly, ‘choice’ experiments - where poor- and rich diets are fed to flies in conjunction – result in heightened mortality, compared to continuous feeding (Ro *et al.*, 2016). These effects are dependent on serotonin signalling (Ro *et al.*, 2016), suggesting that the *perceived*, rather than *actual* composition of food ingested modulates ageing (Libert *et al.*, 2007).

In light of this, it is important to consider the renewed interest in intermittent fasting in both rodent and human studies (Fontana and Partridge, 2015; Mattson, Longo and Harvie, 2017). Studies in the previous century on rodents already demonstrated that inducing intermittent fasting, by feeding animals every other day or by other means, extends lifespan in a similar manner to caloric restriction (reviewed in Anson, Jones and de Cabod, 2005). Two recent studies in mice suggest the same, although effects are not as large as full caloric restriction (Mitchell *et al.*, 2019) and outcomes for systemic ageing have been questioned (Xie *et al.*, 2017). Human data on intermittent fasting is promising (Mattson, Longo and Harvie, 2017) and has potential application in specific diseases (Cignarella *et al.*, 2018), but conclusive evidence from clinical trials is currently lacking (Horne, Muhlestein and Anderson, 2015; Patterson *et al.*, 2015). Our work now suggests that intermittent DR, dependent on its duration, has negative consequences. These observations fit with the ‘refeeding syndrome’ - a clinical condition that occurs at refeeding after a period of starvation (Mehanna, Moledina and Travis, 2008). It remains to be determined which duration of starvation or DR would instigate such harmful physiological effects upon refeeding to the extent that it offsets its physiological benefits in humans.

At present, no mechanistic explanation is apparent which explains the exacerbated mortality when flies return to a rich diet after a period of DR. We have excluded water balance, microbiome, sex-specific and social effects being wholly responsible for our observations. We therefore conclude that in conjunction with physiological costs associated with a rich diet there are hidden costs associated with DR. These costs appear only when a rich diet is resumed after DR. The difference in mortality rates between our switching treatments (Fig. 2B, C, E, F) demonstrate a minimum period of acclimation to a restricted diet is necessary to generate the detectable costs of it. This suggests a physiological change at DR that makes animals more sensitive to rich diets, directly contrary to expectations that follow from evolutionary theory. Drawing from our observation of exacerbated mortality upon resumption of a rich diet - even when DR caused starvation - we suggest the exacerbation of mortality on a rich diet results from physiological adaptations that compensate for the lack of certain components within a restricted diet. We suggest this compensation sensitises animals to the physiological costs associated with the elevated intake, or metabolism of such a specific dietary component, producing exacerbated mortality. For example, such physiological compensation at DR could result, upon resumption of the high nutrient diet, in a higher influx of specific dietary components, or a higher flux in metabolic pathways upregulated due to starvation. Intriguingly these same, otherwise hidden, mechanisms might also underlie why animals fed rich diets continuously are shorter lived than those on DR. This novel paradigm also explains why flies respond rapidly and repeatedly to DR: as an escape from costs associated with the intake or metabolism of a (or several) dietary component(s).

All current evidence so far suggests that uptake of the macronutrient protein is responsible for the effects of diet on longevity (Min *et al.*, 2007; Lee *et al.*, 2008; Maklakov *et al.*, 2008; Solon-Biet *et al.*, 2014; Jensen *et al.*, 2015; Fontana *et al.*, 2016). We suggest that DR’s effect on longevity is not via increased investment in somatic maintenance, but the result from a forced escape from the intrinsically harmful effects of dietary protein. The reason why animals would still choose to eat or absorb intrinsically harmful components, such as protein from their diets, is most likely for its use in reproduction (Speakman and Mitchell, 2011; Jensen *et al.*, 2015). The specific physiological mechanisms that underlie these costs, lie at the heart of DR’s lifespan extending capacities. Our identification of novel dietary phenotypes in the fly that expose these otherwise hidden costs could prove a powerful new experimental phenotype for the mechanistic study of DR. We suggest that the quest to identify the mechanisms of DR will be aided by acceptance that investment in somatic maintenance is not necessarily responsible for the life-extension seen under DR.

## Materials and Methods

### Fly husbandry

Wild-type inbred isofemale flies from the Drosophila melanogaster Genetic Reference Panel (MacKay *et al.*, 2012) were acquired from the Bloomington Stock Centre and the lab of Bart Deplancke (EPFL). Flies were cultured on rich media (8% autolysed yeast, 13% table sugar, 6% cornmeal, 1% agar and nipagin 0.225% [w/v]) with bottles for growing and mating, containing an additional 1% [v/v] propanoic acid. For lifespan experiments, adult flies were subsequently provided with either the same rich media, or a restricted media (2% autolysed yeast) in vials. Restricted media retained the composition of all other media components, given the dietary protein axis is the main lifespan determinant in flies (Lee *et al.*, 2008; Jensen *et al.*, 2015). Cooked fly media was kept for a maximum of 2 weeks at 4-6 °C and was warmed to 25°C before use.

### Experimental mortality protocol and demography cages

Flies were expanded in bottles (Drosophila PP Flask Square Bottom; Flystuff) on a rich diet. Experimental flies were grown in bottles (incubated at 25°C) sprinkled with granulated live yeast, in which 12 females and 2 males had been egg-laying for a period of ∼60 hours. Bottles were sprinkled with water, daily, if media appeared dry until pupation began. Upon eclosion, the adult F1 generation was transferred, daily to generate age-matched cohorts, to mating bottles for 48 hours before being sorted under light CO_2_ anaesthesia (Flystuff Flowbuddy; < 5L / min) (Bartholomew *et al.*, 2015) and transferred to purpose-built demography cages (Good and Tatar, 2001). Lifespan experiments were carried out in a climate-controlled room (12:12 LD, 25°C and 50-60% relative humidity). Cages contained between 100-125 females each; the number of cages was treatment-dependent. All flies were kept on rich media until age 3-6 days whereupon they were divided between the dietary treatments. Individual lifespan was determined from the time when the individual entered the experimental cage (at two days of age) until death or censoring. A census of flies was taken every other day: dead flies were counted and removed, and fresh media was provided at this time. Flies that were alive, but stuck to the side of the vial; escaped flies and individuals affixed to the food (∼10.5% of deaths) were right-censored.

### Fecundity

A subsection of fly feeding vials were imaged and analysed using QuantiFly (Waithe *et al.*, 2015) to determine relative amounts of egg laying.

### Dietary regimes

Two main temporal dietary regimes were imposed on several genotypes of mainly female flies using two diets, restricted (DR, 2% yeast) and rich (8% yeast) with controls of continuous exposure to these diets.

1) To test whether a prolonged period of DR resulted in superior survival and reproduction when conditions improved, flies were exposed to continuous restricted diet that was switched to a rich diet at ∼45-60% survival of the continuous rich diet group (‘long-switch’). All flies of the same genotype were switched on the same day, irrespective of eclosion date.
2) We further tested whether short bouts of DR had similar effects, which also allowed us to test whether effects observed in the long-switch regime were exclusive to older flies. In these diets flies were repeatedly switched between restricted and rich diets at four-day intervals (‘four-day switch’). By starting half of the experimental cohort on restricted or rich diets, current dietary treatments were mirrored and balanced across the cohort.

These experiments were performed on DGRP-195 at high sample size (N = 14,102). Subsequently, to test whether these effects were general, these experiments were expanded to a panel of DGRP lines (DGRP-105; 136; 195; 217; 239; 335; 362; 441; 705; 707; 853) in one large experiment of N = 37,897. Several other parts of the experiments (see below) were run separately (for specific grouping see supplementary data). Dietary treatments were balanced for age. From this experiment, fecundity estimates were also taken from feeding vials, on four consecutive scoring days (for 4-day switch, and continuous treatments) and one scoring day before, and after, the dietary switch (for long-switch, and continuous rich treatment).

### Supplementary dietary regimes

We tested a range of other dietary regimes to test specific hypotheses, alongside the treatments listed above, using line DGRP-195. 1) We tested whether DR could instantly reduce mortality by imposing a short duration (four days) of DR, in late-life, *sensu* Mair *et al.*, 2003, before returning to a rich diet (‘short reverse-switch’). 2) We increased the frequency of the dietary switch to two days (‘2-day switch’) to investigate the length of DR necessary for the observed phenotypes, and 3) further changed the ratio of the time spend on either diet; two days of either rich or restricted diet, to four days of the reverse (‘4-to-2-day switch’).

### Tests of specific hypotheses: microbiome, water balance, sex and social effects

We tested whether the dietary phenotypes observed were due to four potential previously suggested confounding factors: 1) the microbiome (Wong, Dobson and Douglas, 2014), 2) water balance (Fanson, Yap and Taylor, 2012) and 3) social effects (Leech, Sait and Bretman, 2017; Chakraborty *et al.*, 2019). 4) Sex-differences in the DR response (Magwere, Chapman and Partridge, 2004; Regan *et al.*, 2016). These were confirmed not to interfere with the observed phenotype (see results). DGRP-195 were used exclusively for these experiments, under the continuous restricted, continuous rich, and long-switch diets. 1) We assessed whether disruption of the gut microbiome was responsible for the mortality phenotype observed by wholesale abating the microbiome. Flies were provided media upon which an array of broad-spectrum antibiotics (50 μl of a stock solution, comprised of 100 μg/ml ampicillin, 50 μg/ml vancomycin, 100 μg/ml neomycin, and 100 μg/ml metronidazole) were pipetted and left for 24hrs. We assumed dissolution incorporation in the top 1 ml of food (Ren, Finkel and Tower, 2009). Antibiotic treatment began four days prior to dietary switch treatments, and concluded eight days thereafter. Ablation of the microbiome was confirmed by whole-fly homogenisation (age 20 days; 8 days post-antibiotic treatment) and growth of solution on MRS agar plates (Oxoid; see Fig. S7). Individuals (6 control and 6 antibiotic treated) were removed from cages containing a continuous restricted diet, washed in ethanol, and rinsed in PBS (Gibco). Homogenisation took place in 500 μl of PBS, and solute was transferred to a 96-well plate for 1:10 serial dilutions. Dilutions were spotted on plates with, and without antibiotic (500 μl of stock solution) and incubated at 25°C for 72 hours. Plates were coated with parafilm to mimic anoxic conditions. 2) Flies were provided with ∼1cm^3^ portion of water-agar (2%[w/v]) accompanying media in vials, to eliminate desiccation as a proximal cause. Water-agar supplementation began at age four and continued throughout the flies’ full life course. 3) Social effects were excluded by housing flies individually in vials. These flies were taken from experimental cages and put on the experimental diets at the dietary switch 4) Males were assessed for mortality in the 4 day switch dietary regime.

### Experimental batches

All demography experiments contained the relevant controls, grown and assayed for mortality at the same time. Where data are plotted in a single figure, this constitutes results gathered from a batch of flies at the same chronological time.

### Data Analysis

Mixed cox-proportional hazard models were used that included ‘cage’ as random term to correct for uncertainty of pseudo-replicated effects within demography cages (Ripatti and Palmgren, 2000; Therneau *et al.*, 2003). We used interval-based models that used time-dependent covariates to estimate the differential mortality risks associated with diet (and with time spend on a diet - after diets changed), as imposed in the different dietary regimes. These models allow a statistical association, within the cox-proportional hazard risk, with the current state (e.g. diet) and mortality. Flies in the long-switch dietary regime were also analysed in a state-dependent manner, coding for long-switch only when this state change occurred. Repeated switching regimes were considered lifelong treatments and tested in interaction with the state variable diet. Each model used continuous rich food and DGRP-195 as reference category, except if otherwise stated.

Interactions between dietary regime, diet and genotype were fitted to test for differential effects of diet on mortality depending on the regime it was provided. Additional specific tests of coefficients are provided that combine the single and interaction term (in a z-test, using the maximum s.e. of the factor compared) to test how mortality risk was changing compared to specific reference categories of interest (e.g. compared to continuous DR). For comparisons between genotypes we report full models including all data and models fitted within each genotype separately. The latter corrects for deviations in proportionality of hazards between the genotypes. Qualitative conclusions remain similar, and formal tests for proportionality of hazards are not available for mixed effects cox regressions. Models without a time-dependent covariate for diet were also run to compare overall longevity differences as a result of alternating exposure to DR (2-day switch, 4-day switch and their combination). These models therefore test the integrated effect on mortality disregarding any within dietary treatment diet effects. Coefficients are reported as logged hazards with significance based on z-tests. Right-censoring was included, as indicated above.

Egg laying was analysed as a mixed generalized Poisson model using cage as random term, and fitting age as a non-continuous factor in the analysis. Estimates from models are presented (effects of dietary regime) as well as model comparisons using log-likelihood comparison with chi-square to test overall effects of genotype. Effects of different dietary regimes were estimated within the same model. Comparisons of genotypic effects were performed for each different dietary regime separately compared to continuous treatment, as not to conflate genetic variance across different categories with each other.

## Acknowledgements

AWM is supported by the NERC ACCE Doctoral Training Program. MT is supported by American Federation of Aging Research, Grant/Award Number: GR5290420; National Institute on Aging, Grant/Award Number:R37 AG024360, T32 AG 41688. MJPS is supported by a Sir Henry Wellcome and a Sheffield Vice Chancellor’s Fellowship, and the Natural Environment Research Council (M005941 & N013832). We thank Laura Carrilero and James PJ Hall from Michael Brockhurst’s lab for their valuable assistance during the microbiome ablation experiment. We also thank Kang-Wook Kim for support sorting flies and the Deplancke lab for supplying DGRP lines.

## Supplementary figures

**Figure S1.**
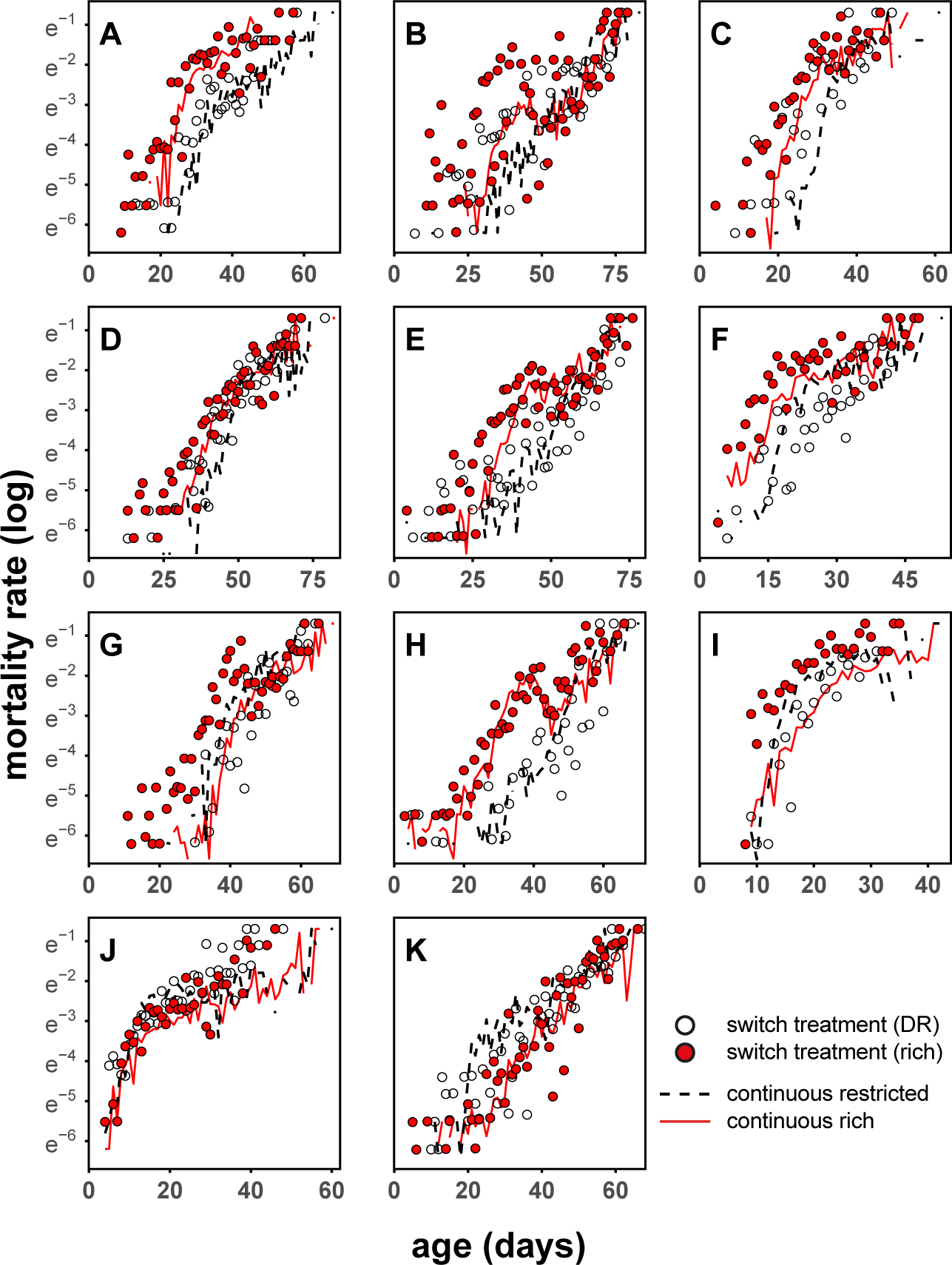
4-day switch treatment in a panel of 11 DGRP genotypes. A – 195; B – 105; C – 217; D – 441; E – 705; F – 707; G – 136; H – 362; I – 239; J – 335; K – 853. Continuous rich, and restricted treatments plotted as lines (solid red and dashed black, respectively). Switch treatments plotted as points (white and red). The exacerbation of mortality due to switch phenotypes is observable as the difference between mortality at continuous rich diet (red line), and mortality of switch treatment when on a rich diet (red points). N = 29,740 total; ∼2,725 per genotype; 13,375 for continuous rich treatments, and ∼8,170 for continuous rich and 4-day switch treatments. Dietary switch for 4-day switch treatment group occurred every 4 days, and was mirrored at each time point. Continuous rich and restricted treatments are twinned with long switch treatment experiment (Fig. 2). All panels contain daily time-points, as in Fig.2.

**Figure S2.**
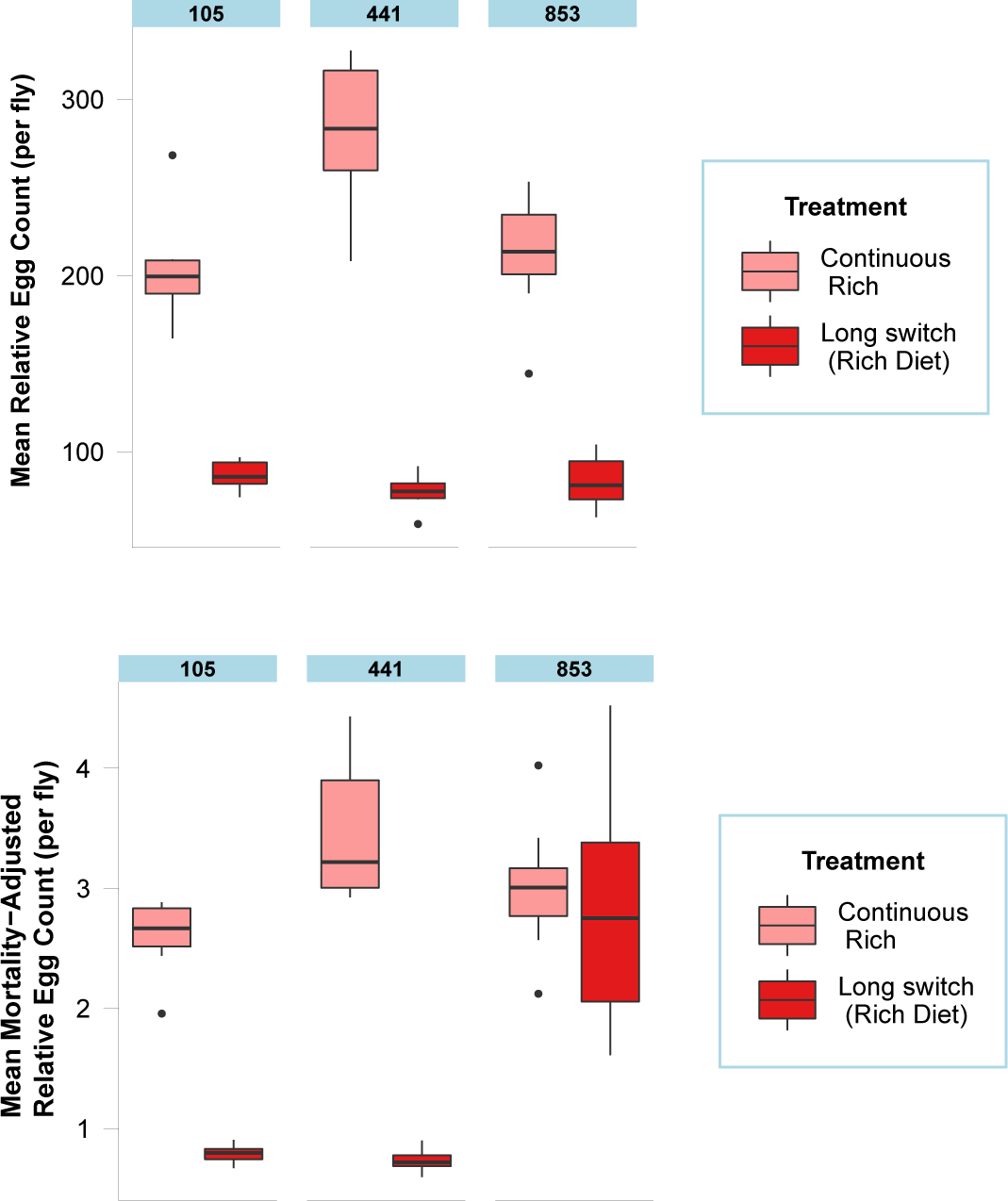
Fecundity analysis of long switch treatment from 3 DGRP genotypes. No compensation via fecundity for reduced lifespans in switch treatment. Raw (above) and mortality corrected (below) egg counts of DGRP-105; 441; 853 from long switch treatment experiment (Fig. 2). Counts generated using QuantiFly software. Counts are relative, but directly comparable. Flies assayed between age 44-47 days, with boxplots (median, with the box depicting a quartile each way, and whiskers showing the range; outliers plotted as dots) aggregating totals. Each cage was assayed once, on the first scoring day post dietary switch. Mortality corrected counts (below) generated by dividing raw counts, by N flies remaining in cage at the time of assaying. N = on average, 7 cages assayed, per treatment, per genotype.

**Figure S3.**
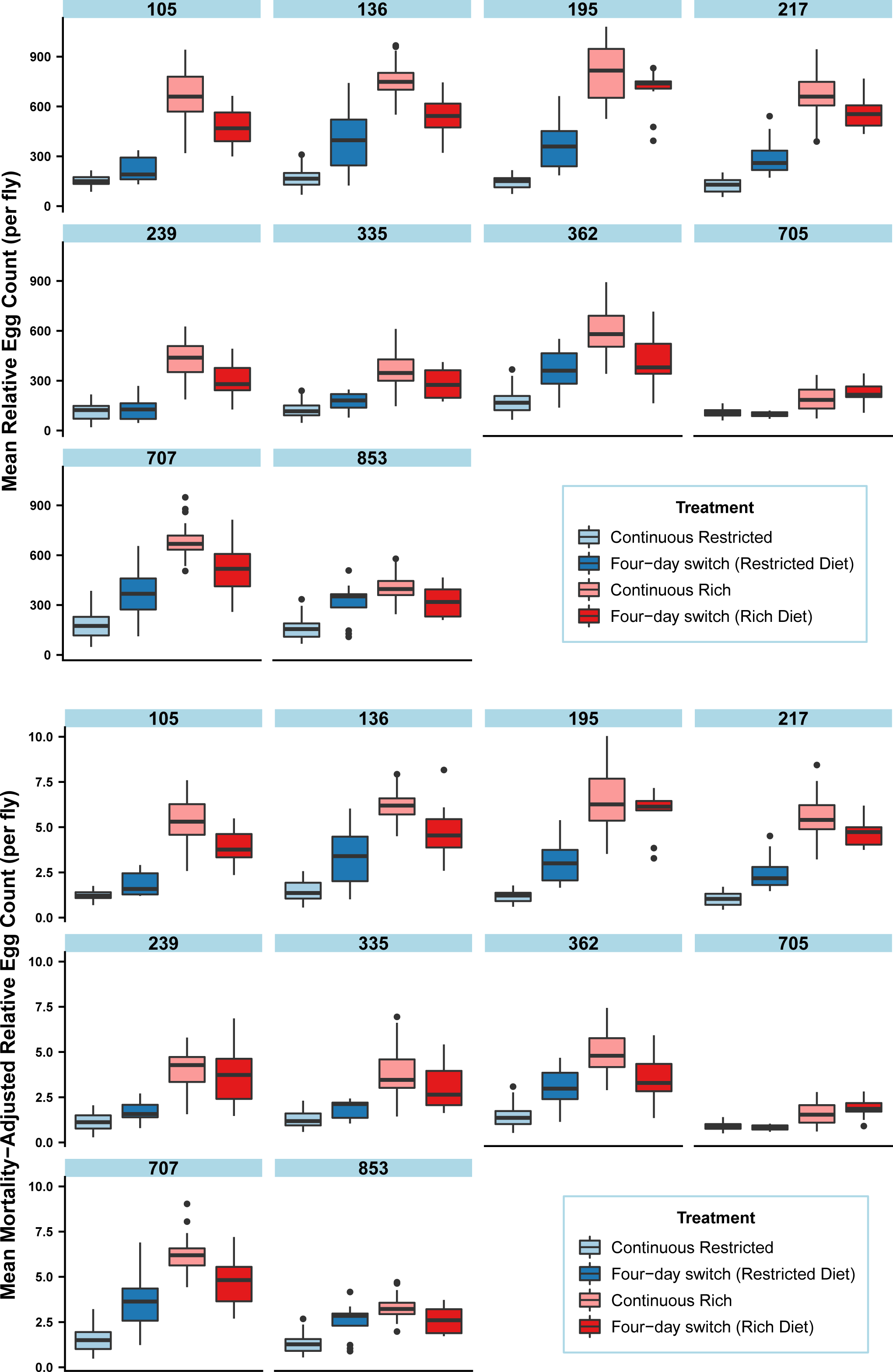
Fecundity analysis of 4-day switch treatment from 10 DGRP genotypes. No compensation via fecundity for reduced lifespans in switch treatment. Raw (above) and mortality corrected (below) egg counts of DGRP-105; 136; 195; 217; 239; 335; 362; 705; 707; 853 from long switch treatment experiment (Fig. 2). Counts generated using QuantiFly software. Counts are relative, but directly comparable. Flies assayed between age 8-21 days, with boxplots aggregating totals (median, with the box depicting a quartile each way, and whiskers showing the range; outliers plotted as dots). Each cage was assayed on 4 consecutive scoring days. Mortality corrected counts (below) generated by dividing raw counts, by N flies remaining in cage at the time of assaying. N = on average, 7 cages assayed, per treatment, per genotype.

**Figure S4.**
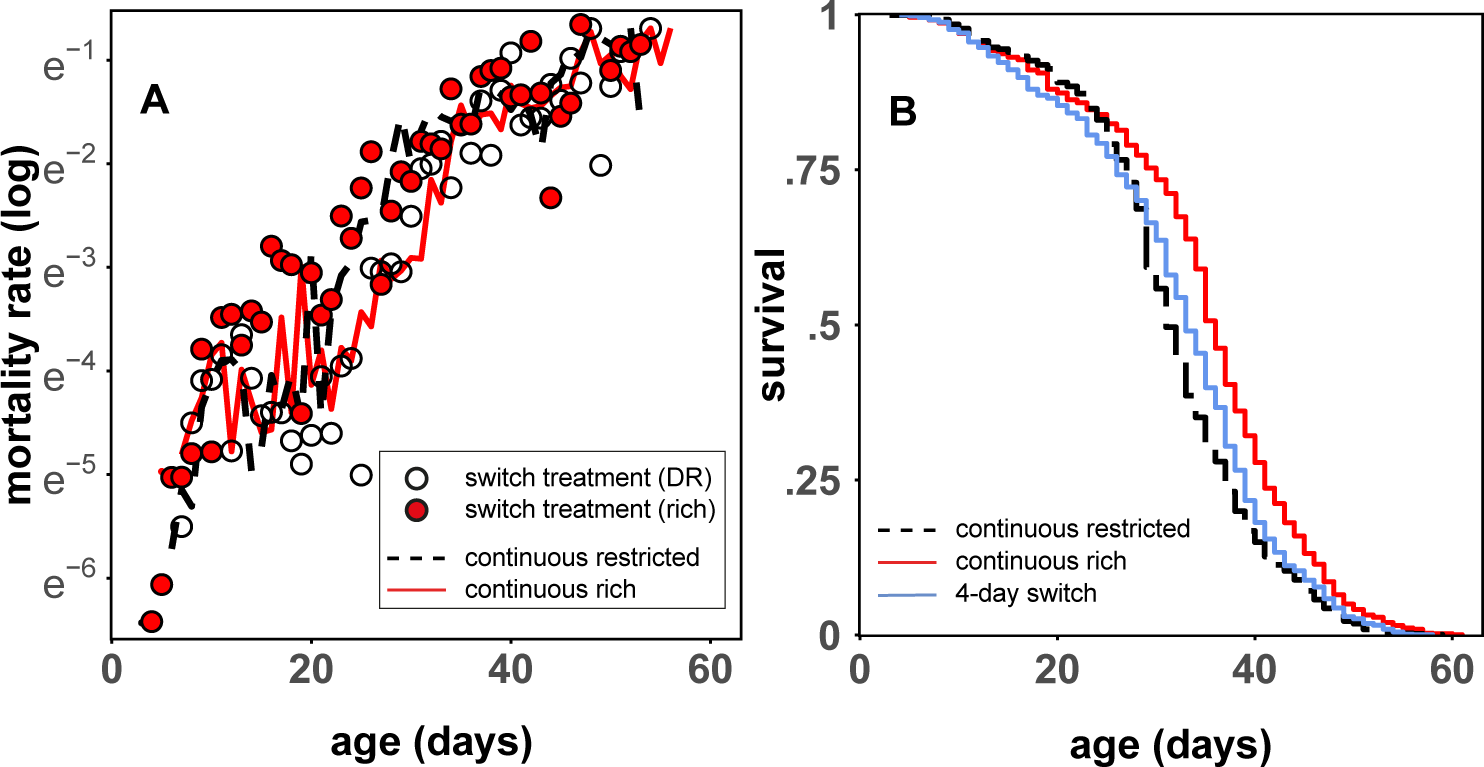
4-day switch treatment of DGRP-195 males. A – 4-day switch mortality; B – 4-day switch survival. Muted response to 4-day switch treatment in males. Rich diet in the 4-day switch increased mortality compared to continuously rich fed flies. Continuous rich, and restricted treatments plotted as lines (solid red and dashed black, respectively). Switch treatment plotted as points (white and red). The exacerbation of mortality due to switch phenotypes is observable as the difference between mortality at continuous rich diet (red line), and mortality of switch treatment when on a rich diet (red points). N = 4,429 total; ∼1,475 per treatment. Dietary switch for 4-day switch treatment group occurred every 4 days, and was mirrored at each time point. Both panels contain daily time-points, as in Fig.2.

**Figure S5.**
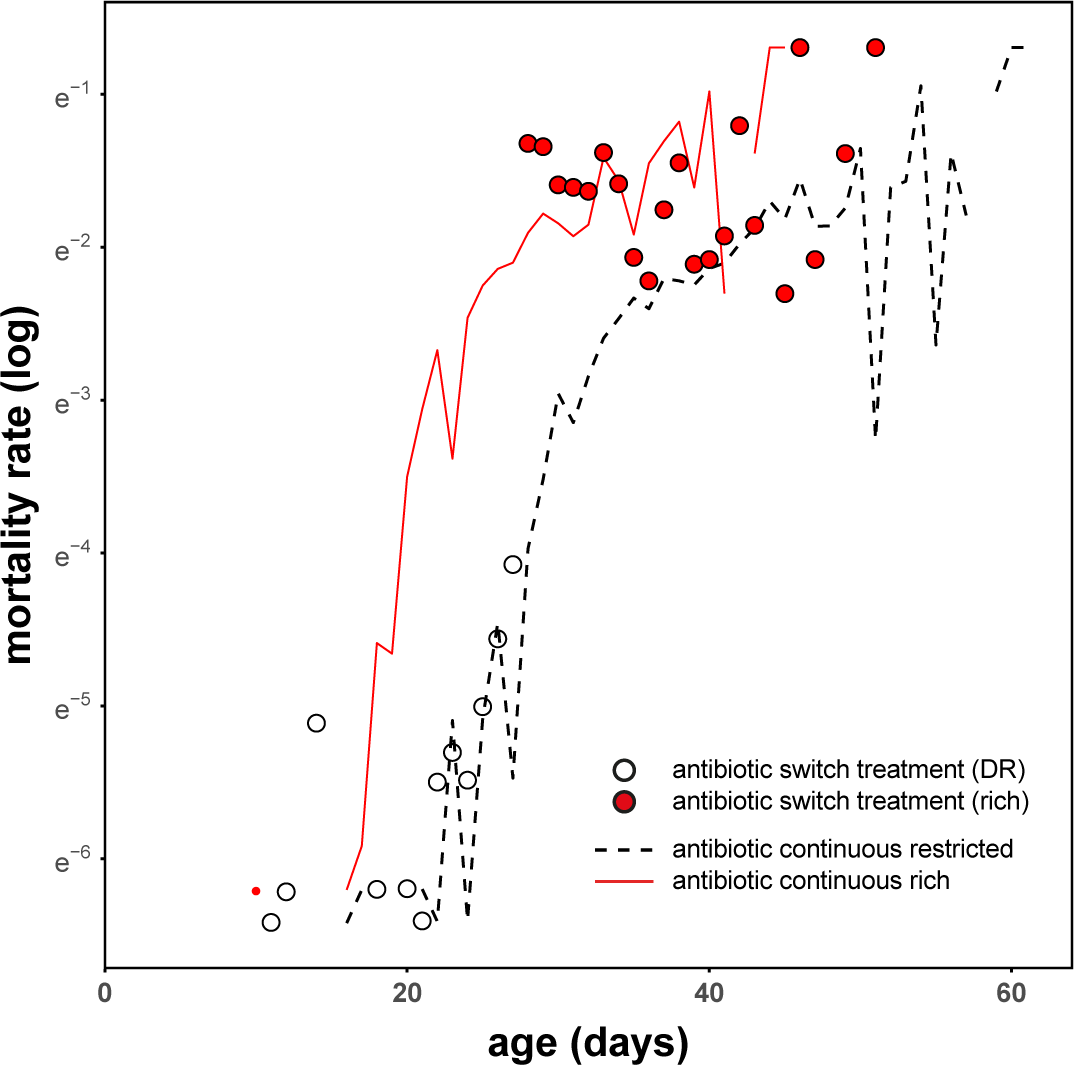
Antibiotic long switch treatment of DGRP-195. Long switch phenotype independent of antibiotic treatment. Antibiotic treatment took place in all treatments four days prior to dietary switch, and concluded eight days thereafter. Continuous rich, and restricted treatments plotted as lines (solid red and dashed black, respectively). Switch treatment plotted as points (white and red). The exacerbation of mortality due to switch phenotypes is observable as the difference between mortality at continuous rich diet (red line), and mortality of switch treatment when on a rich diet (red points). N = 2,605 total; ∼870 per treatment. (See Fig. S7 for confirmation of ablation of microbiome). Figure contains daily time-points, as in Fig.2.

**Figure S6.**
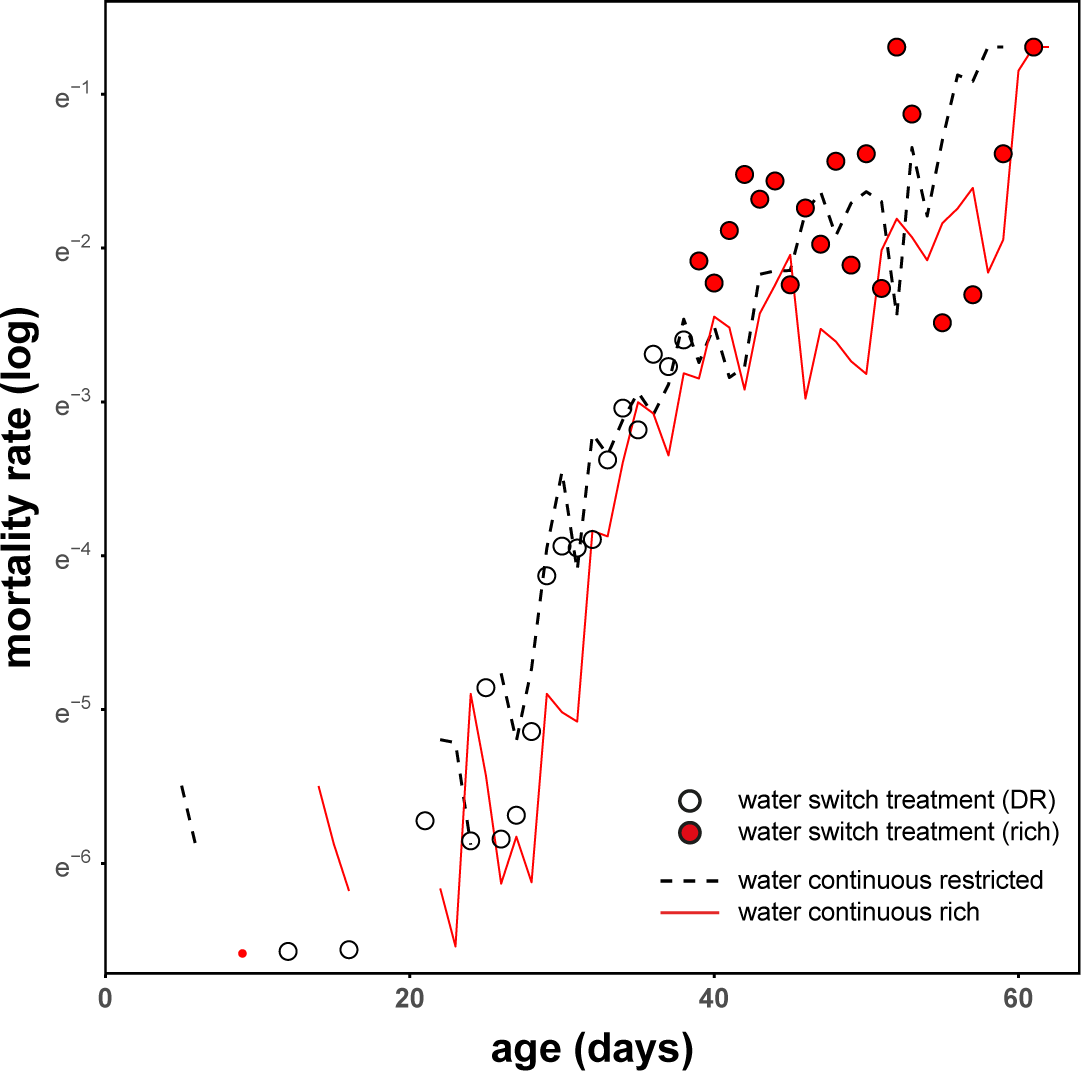
Water supplemented long switch treatment of DGRP-195. Long switch phenotype independent of water supplementation. Water supplementation took place in all treatments throughout life of the cage. Continuous rich, and restricted treatments plotted as lines (solid red and dashed black, respectively). Switch treatment plotted as points (white and red). The exacerbation of mortality due to switch phenotypes is observable as the difference between mortality at continuous rich diet (red line), and mortality of switch treatment when on a rich diet (red points). N = 2,562 total; ∼850 per treatment. **NB** water supplementation did change the response to DR. This effect was followed up with an experiment containing five different genotypes across a range of diets, with only a shift in reaction norm detected (manuscript in preparation). DR is not explained by dehydration, as is sometimes suggested, nor is the long switch phenotype. Figure contains daily time-points, as in Fig.2.

**Figure S7/Table S7.**
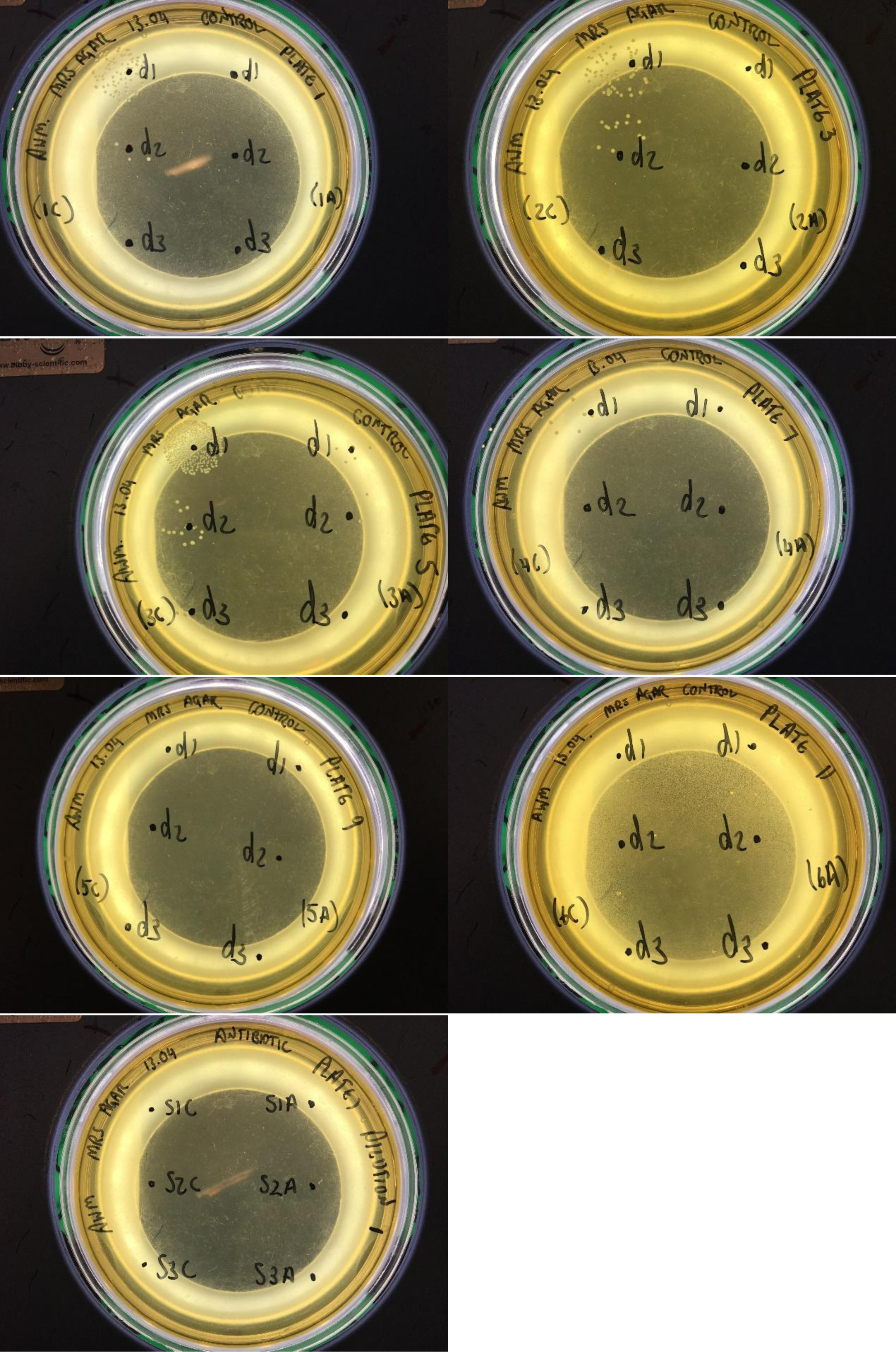
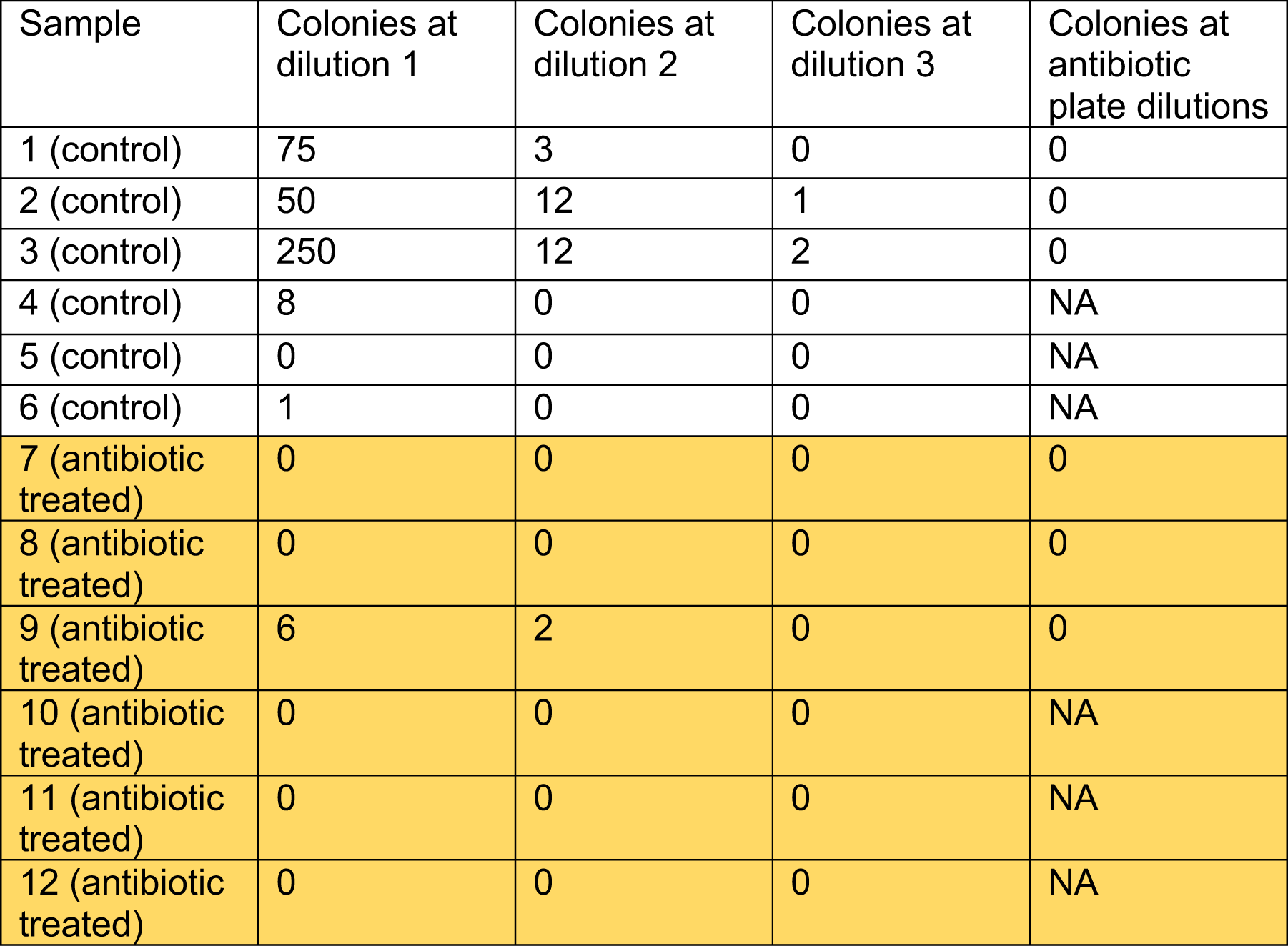
Confirmation of ablation of microbiome. Images of bacterial colonies visible on MRS agar plates (above) and estimated colony count (below). Twelve samples ceded from control, or antibiotic-treated cages. Lysate was diluted post-homogenisation and grown on control, or antibiotic-treated plates. No growth visible under antibiotic treated plate conditions. 98.4% reduction of total microbiota observed at dilution 1. 92.5% reduction of total microbiota observed at dilution 2.

**Table S1.** Effect of dietary regimes on interval-based log hazard ratios of mortality in DGRP-195.

| coefficient | Full Model |  |  |  | Versus continuous DR |  |  |
| --- | --- | --- | --- | --- | --- | --- | --- |
|  | estimate | exp (estimate) | s.e. | p | estimate | exp (estimate) | p |
| DR | -1.179 | 0.308 | 0.047 | **<0.001** |  |  |  |
| long switch | 1.310 | 3.705 | 0.055 | **<0.001** |  |  |  |
| 4-day switch | 0.858 | 2.358 | 0.062 | **<0.001** |  |  |  |
| 2-day switch | 0.125 | 1.133 | 0.063 | **0.047** |  |  |  |
| short reverse-switch | 0.272 | 1.312 | 0.051 | **<0.001** |  |  |  |
| 4-day switch * DR | -0.409 | 0.665 | 0.085 | **<0.001** | 0.449 | 1.567 | **<0.001** |
| 2-day switch * DR | 0.255 | 1.291 | 0.074 | **0.001** | 0.380 | 1.462 | **<0.001** |
| short switch * DR | -0.466 | 0.627 | 0.116 | **<0.001** | -0.195 | 0.823 | 0.092 |

**Table S2.**
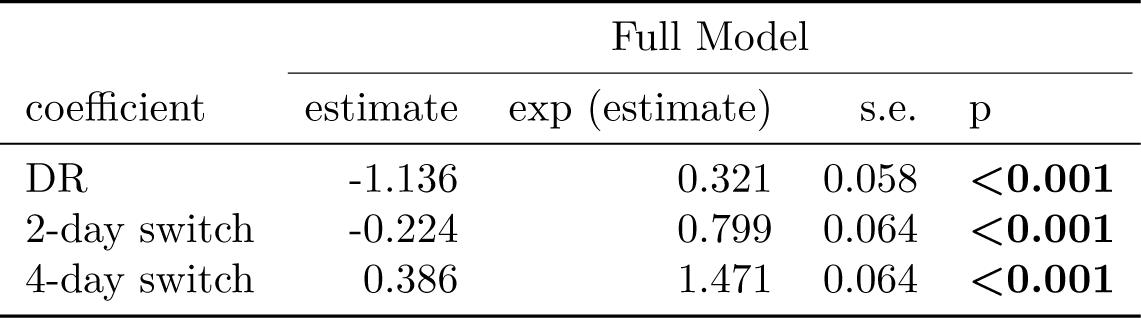
Effect of dietary regimes on longevity in DGRP-195.

**Table S3.** Time-dependent effect of mortality increase induced by a long-switch from reduced to rich diets in DGRP-195

| coefficient | Full Model |  |  |  |
| --- | --- | --- | --- | --- |
|  | estimate | exp (estimate) | s.e. | p |
| day 2 | 1.629 | 5.101 | 0.085 | **<0.001** |
| day 4 | 0.847 | 2.334 | 0.093 | **<0.001** |
| day 6 | 0.452 | 1.572 | 0.121 | **<0.001** |
| day 8 | -0.045 | 0.956 | 0.216 | 0.84 |
| day 10 | -0.226 | 0.798 | 0.369 | 0.54 |
| day 12 | -0.077 | 0.926 | 0.601 | 0.9 |
| > day 14 | -1.038 | 0.354 | 1.066 | 0.33 |

**Table S4.** Time-dependent effect of mortality increase induced by a 4-day switch in DGRP-195

| coefficient | Full Model |  |  |  |
| --- | --- | --- | --- | --- |
|  | estimate | exp (estimate) | s.e. | p |
| DR | -1.303 | 0.272 | 0.094 | **<0.001** |
| 2nd interval | 0.227 | 1.255 | 0.055 | **<0.001** |
| 2nd interval * DR | -0.715 | 0.489 | 0.143 | **<0.001** |

**Table S5.** Effect of asymmetrical dietary regimes on mortality in DGRP-195.

| coefficient | Full Model |  |  |  | Versus continuous DR |  |  |
| --- | --- | --- | --- | --- | --- | --- | --- |
|  | estimate | exp (estimate) | s.e. | p | estimate | exp (estimate) | p |
| DR | -1.684 | 0.186 | 0.087 | **<0.001** |  |  |  |
| 4d-DR\|2d-Rich | -0.209 | 0.811 | 0.097 | **0.032** |  |  |  |
| 2d-DR\|4d-Rich | -0.157 | 0.854 | 0.095 | 0.097 |  |  |  |
| 4d-DR\|2d-Rich * DR | 0.458 | 1.581 | 0.113 | **<0.001** | 0.249 | 1.283 | **0.027** |
| 2d-DR\|4d-Rich * DR | 0.929 | 2.531 | 0.124 | **<0.001** | 0.771 | 2.163 | **<0.001** |

**Table S6.** Effect of asymmetrical dietary regimes on longevity in DGRP-195.

| coefficient | Full Model |  |  |  |
| --- | --- | --- | --- | --- |
|  | estimate | exp (estimate) | s.e. | p |
| DR | -1.692 | 0.184 | 0.087 | **<0.001** |
| 4d-DR\|2d-Rich | -0.805 | 0.447 | 0.095 | **<0.001** |
| 2d-DR\|4d-Rich | -0.343 | 0.710 | 0.094 | **<0.001** |

**Table S7.** Mortality increases in response to a rich diet after a period of DR (long-switch) across a panel of 11 DGRP lines (195 is reference)

| coefficient | Full Model |  |  |  | Effect of DR |  |  | Long switch versus rich diet |  |  |
| --- | --- | --- | --- | --- | --- | --- | --- | --- | --- | --- |
|  | estimate | exp | s.e. | p | estimate | exp | p | estimate | exp | p |
| DR | -1.723 | 0.179 | 0.157 | **<0.001** |  |  |  |  |  |  |
| long switch | 0.973 | 2.645 | 0.160 | **<0.001** |  |  |  |  |  |  |
| 105 | -2.681 | 0.068 | 0.146 | **<0.001** |  |  |  |  |  |  |
| 136 | -1.969 | 0.140 | 0.152 | **<0.001** |  |  |  |  |  |  |
| 217 | -0.080 | 0.923 | 0.159 | 0.61 |  |  |  |  |  |  |
| 239 | 0.899 | 2.457 | 0.156 | **<0.001** |  |  |  |  |  |  |
| 335 | 0.190 | 1.209 | 0.155 | 0.22 |  |  |  |  |  |  |
| 362 | -1.022 | 0.360 | 0.155 | **<0.001** |  |  |  |  |  |  |
| 441 | -2.107 | 0.122 | 0.152 | **<0.001** |  |  |  |  |  |  |
| 705 | -1.861 | 0.156 | 0.154 | **<0.001** |  |  |  |  |  |  |
| 707 | 1.073 | 2.925 | 0.156 | **<0.001** |  |  |  |  |  |  |
| 853 | -1.803 | 0.165 | 0.152 | **<0.001** |  |  |  |  |  |  |
| 105 * DR | 1.296 | 3.655 | 0.217 | **<0.001** | -0.427 | 0.653 | **0.049** |  |  |  |
| 136 * DR | 1.995 | 7.355 | 0.213 | **<0.001** | 0.273 | 1.314 | 0.199 |  |  |  |
| 217 * DR | 0.622 | 1.863 | 0.219 | **0.004** | -1.100 | 0.333 | **<0.001** |  |  |  |
| 239 * DR | 2.427 | 11.319 | 0.211 | **<0.001** | 0.704 | 2.022 | **0.001** |  |  |  |
| 335 * DR | 2.947 | 19.052 | 0.208 | **<0.001** | 1.225 | 3.403 | **<0.001** |  |  |  |
| 362 * DR | 0.343 | 1.409 | 0.219 | 0.12 | -1.379 | 0.252 | **<0.001** |  |  |  |
| 441 * DR | 1.002 | 2.724 | 0.215 | **<0.001** | -0.720 | 0.487 | **0.001** |  |  |  |
| 705 * DR | 0.602 | 1.826 | 0.217 | **0.006** | -1.120 | 0.326 | **<0.001** |  |  |  |
| 707 * DR | 0.886 | 2.424 | 0.215 | **<0.001** | -0.837 | 0.433 | **<0.001** |  |  |  |
| 853 * DR | 2.549 | 12.796 | 0.210 | **<0.001** | 0.827 | 2.286 | **<0.001** |  |  |  |
| 105 * long switch | 0.168 | 1.183 | 0.220 | 0.44 |  |  |  | 1.141 | 3.130 | **<0.001** |
| 136 * long switch | -0.381 | 0.683 | 0.222 | 0.086 |  |  |  | 0.592 | 1.807 | **0.008** |
| 217 * long switch | 1.152 | 3.165 | 0.226 | **<0.001** |  |  |  | 2.125 | 8.370 | **<0.001** |
| 239 * long switch | 0.870 | 2.387 | 0.224 | **<0.001** |  |  |  | 1.843 | 6.313 | **<0.001** |
| 335 * long switch | -0.305 | 0.737 | 0.221 | 0.17 |  |  |  | 0.668 | 1.950 | **0.002** |
| 362 * long switch | -0.982 | 0.375 | 0.223 | **<0.001** |  |  |  | -0.009 | 0.991 | 0.968 |
| 441 * long switch | -0.172 | 0.842 | 0.220 | 0.43 |  |  |  | 0.800 | 2.226 | **<0.001** |
| 705 * long switch | -0.605 | 0.546 | 0.222 | **0.006** |  |  |  | 0.368 | 1.445 | 0.097 |
| 707 * long switch | 0.188 | 1.206 | 0.218 | 0.39 |  |  |  | 1.160 | 3.191 | **<0.001** |
| 853 * long switch | 0.117 | 1.124 | 0.225 | 0.6 |  |  |  | 1.090 | 2.973 | **<0.001** |

**Table S8.** Models run within each genotype testing for increases in response to a rich diet after a period of DR (long-switch)

| coefficient | Estimates from individual models |  |  |  |
| --- | --- | --- | --- | --- |
|  | estimate | exp | s.e. | p |
| 105 DR | -0.501 | 0.149 | 0.606 | **0.001** |
| 136 DR | 0.647 | 0.120 | 1.910 | **<0.001** |
| 195 DR | -1.737 | 0.197 | 0.176 | **<0.001** |
| 217 DR | -1.102 | 0.178 | 0.332 | **<0.001** |
| 239 DR | 0.775 | 0.071 | 2.170 | **<0.001** |
| 335 DR | 0.570 | 0.115 | 1.768 | **<0.001** |
| 362 DR | -1.201 | 0.130 | 0.301 | **<0.001** |
| 441 DR | -0.503 | 0.101 | 0.605 | **<0.001** |
| 705 DR | -0.935 | 0.095 | 0.393 | **<0.001** |
| 707 DR | -0.646 | 0.076 | 0.524 | **<0.001** |
| 853 DR | 0.828 | 0.093 | 2.288 | **<0.001** |
| 105 long switch | 1.258 | 0.156 | 3.520 | **<0.001** |
| 136 long switch | 0.113 | 0.132 | 1.120 | 0.39 |
| 195 long switch | 0.453 | 0.203 | 1.573 | **0.025** |
| 217 long switch | 1.650 | 0.187 | 5.205 | **<0.001** |
| 239 long switch | 1.483 | 0.103 | 4.406 | **<0.001** |
| 335 long switch | 0.881 | 0.136 | 2.413 | **<0.001** |
| 362 long switch | 0.145 | 0.137 | 1.157 | 0.29 |
| 441 long switch | 0.313 | 0.109 | 1.368 | **0.004** |
| 705 long switch | 0.329 | 0.102 | 1.389 | **0.001** |
| 707 long switch | 0.854 | 0.083 | 2.348 | **<0.001** |
| 853 long switch | 1.373 | 0.122 | 3.949 | **<0.001** |

**Table S9.** Effect of alterating DR and rich diets every 4 days (4-day switch) on longevity across 11 DGRP lines (195 is reference)

| coefficient | Full Model |  |  |  | Effect compared to rich diet |  |  |
| --- | --- | --- | --- | --- | --- | --- | --- |
|  | estimate | exp | s.e. | p | estimate | exp | p |
| 4-day switch | -0.140 | 0.869 | 0.114 | 0.22 |  |  |  |
| 105 | -2.476 | 0.084 | 0.096 | **<0.001** |  |  |  |
| 136 | -1.858 | 0.156 | 0.098 | **<0.001** |  |  |  |
| 217 | -0.065 | 0.937 | 0.100 | 0.52 |  |  |  |
| 239 | 0.918 | 2.505 | 0.099 | **<0.001** |  |  |  |
| 335 | 0.226 | 1.254 | 0.100 | **0.024** |  |  |  |
| 362 | -0.988 | 0.373 | 0.099 | **<0.001** |  |  |  |
| 441 | -1.969 | 0.140 | 0.097 | **<0.001** |  |  |  |
| 705 | -1.769 | 0.171 | 0.098 | **<0.001** |  |  |  |
| 707 | 1.054 | 2.869 | 0.100 | **<0.001** |  |  |  |
| 853 | -1.697 | 0.183 | 0.098 | **<0.001** |  |  |  |
| 105 * 4-day switch | 0.644 | 1.904 | 0.159 | **<0.001** | 0.504 | 1.656 | **0.002** |
| 136 * 4-day switch | 0.632 | 1.880 | 0.157 | **<0.001** | 0.492 | 1.635 | **0.002** |
| 217 * 4-day switch | 0.490 | 1.632 | 0.158 | **0.002** | 0.350 | 1.419 | **0.027** |
| 239 * 4-day switch | 1.053 | 2.866 | 0.157 | **<0.001** | 0.913 | 2.491 | **<0.001** |
| 335 * 4-day switch | 1.026 | 2.789 | 0.158 | **<0.001** | 0.886 | 2.425 | **<0.001** |
| 362 * 4-day switch | -0.343 | 0.710 | 0.157 | **0.029** | -0.483 | 0.617 | **0.002** |
| 441 * 4-day switch | 0.163 | 1.177 | 0.158 | 0.3 | 0.023 | 1.023 | 0.885 |
| 705 * 4-day switch | -0.032 | 0.968 | 0.157 | 0.84 | -0.172 | 0.842 | 0.273 |
| 707 * 4-day switch | 0.153 | 1.165 | 0.158 | 0.33 | 0.013 | 1.013 | 0.935 |
| 853 * 4-day switch | 0.365 | 1.441 | 0.157 | **0.02** | 0.225 | 1.253 | 0.152 |

**Table S10.** Effect of alterating DR and rich diets every 4 days (4 day switch) on mortality at each diet, across 11 DGRP lines (195 is reference)

| coefficient | Full Model |  |  |  | Effect of DR |  |  | Compared to continuous diets |  |  |  |  |  |
| --- | --- | --- | --- | --- | --- | --- | --- | --- | --- | --- | --- | --- | --- |
|  |  |  |  |  |  |  |  | 4 day switch at rich diet |  |  | 4 day switch at DR |  |  |
|  | estimate | exp | s.e. | p | estimate | exp | p | estimate | exp | p | estimate | exp | p |
| DR | -1.378 | 0.252 | 0.129 | **<0.001** |  |  |  |  |  |  |  |  |  |
| 4-day switch | 0.305 | 1.357 | 0.132 | **0.02** |  |  |  |  |  |  |  |  |  |
| 4-day switch * DR | -0.078 | 0.925 | 0.164 | 0.63 |  |  |  |  |  |  |  |  |  |
| 105 | -2.649 | 0.071 | 0.109 | **<0.001** |  |  |  |  |  |  |  |  |  |
| 136 | -1.907 | 0.148 | 0.111 | **<0.001** |  |  |  |  |  |  |  |  |  |
| 217 | -0.064 | 0.938 | 0.115 | 0.58 |  |  |  |  |  |  |  |  |  |
| 239 | 0.907 | 2.476 | 0.113 | **<0.001** |  |  |  |  |  |  |  |  |  |
| 335 | 0.227 | 1.255 | 0.114 | **0.046** |  |  |  |  |  |  |  |  |  |
| 362 | -1.024 | 0.359 | 0.113 | **<0.001** |  |  |  |  |  |  |  |  |  |
| 441 | -2.046 | 0.129 | 0.111 | **<0.001** |  |  |  |  |  |  |  |  |  |
| 705 | -1.846 | 0.158 | 0.112 | **<0.001** |  |  |  |  |  |  |  |  |  |
| 707 | 1.051 | 2.859 | 0.113 | **<0.001** |  |  |  |  |  |  |  |  |  |
| 853 | -1.733 | 0.177 | 0.111 | **<0.001** |  |  |  |  |  |  |  |  |  |
| 105 * DR | 1.171 | 3.224 | 0.180 | **<0.001** | -0.207 | 0.813 | 0.248 |  |  |  |  |  |  |
| 136 * DR | 1.804 | 6.071 | 0.178 | **<0.001** | 0.426 | 1.531 | **0.017** |  |  |  |  |  |  |
| 217 * DR | 0.822 | 2.275 | 0.178 | **<0.001** | -0.556 | 0.574 | **0.002** |  |  |  |  |  |  |
| 239 * DR | 2.046 | 7.740 | 0.179 | **<0.001** | 0.668 | 1.951 | **<0.001** |  |  |  |  |  |  |
| 335 * DR | 2.104 | 8.197 | 0.178 | **<0.001** | 0.726 | 2.066 | **<0.001** |  |  |  |  |  |  |
| 362 * DR | 0.307 | 1.359 | 0.179 | 0.087 | -1.071 | 0.343 | **<0.001** |  |  |  |  |  |  |
| 441 * DR | 0.994 | 2.701 | 0.179 | **<0.001** | -0.384 | 0.681 | **0.031** |  |  |  |  |  |  |
| 705 * DR | 0.527 | 1.695 | 0.178 | **0.003** | -0.851 | 0.427 | **<0.001** |  |  |  |  |  |  |
| 707 * DR | 0.797 | 2.218 | 0.179 | **<0.001** | -0.581 | 0.559 | **0.001** |  |  |  |  |  |  |
| 853 * DR | 2.090 | 8.086 | 0.178 | **<0.001** | 0.712 | 2.038 | **<0.001** |  |  |  |  |  |  |
| 105 * 4-day switch | 0.530 | 1.699 | 0.184 | **0.004** |  |  |  | 0.835 | 2.305 | **<0.001** |  |  |  |
| 136 * 4-day switch | 0.648 | 1.912 | 0.182 | **<0.001** |  |  |  | 0.953 | 2.595 | **<0.001** |  |  |  |
| 217 * 4-day switch | 0.372 | 1.451 | 0.183 | **0.042** |  |  |  | 0.677 | 1.969 | **<0.001** |  |  |  |
| 239 * 4-day switch | 1.019 | 2.771 | 0.182 | **<0.001** |  |  |  | 1.324 | 3.759 | **<0.001** |  |  |  |
| 335 * 4-day switch | 0.451 | 1.569 | 0.186 | **0.015** |  |  |  | 0.756 | 2.129 | **<0.001** |  |  |  |
| 362 * 4-day switch | -0.285 | 0.752 | 0.182 | 0.12 |  |  |  | 0.020 | 1.020 | 0.914 |  |  |  |
| 441 * 4-day switch | -0.158 | 0.854 | 0.184 | 0.39 |  |  |  | 0.148 | 1.159 | 0.421 |  |  |  |
| 705 * 4-day switch | -0.185 | 0.831 | 0.182 | 0.31 |  |  |  | 0.120 | 1.127 | 0.511 |  |  |  |
| 707 * 4-day switch | 0.167 | 1.182 | 0.183 | 0.36 |  |  |  | 0.472 | 1.603 | **0.01** |  |  |  |
| 853 * 4-day switch | -0.140 | 0.869 | 0.185 | 0.45 |  |  |  | 0.165 | 1.179 | 0.372 |  |  |  |
| 105 * 4-day switch * DR | -0.580 | 0.560 | 0.223 | **0.009** |  |  |  |  |  |  | 0.177 | 1.194 | 0.427 |
| 136 * 4-day switch * DR | -1.735 | 0.176 | 0.226 | **<0.001** |  |  |  |  |  |  | -0.859 | 0.423 | **<0.001** |
| 217 * 4-day switch * DR | -0.272 | 0.762 | 0.223 | 0.22 |  |  |  |  |  |  | 0.327 | 1.387 | 0.142 |
| 239 * 4-day switch * DR | -1.846 | 0.158 | 0.225 | **<0.001** |  |  |  |  |  |  | -0.600 | 0.549 | **0.008** |
| 335 * 4-day switch * DR | -0.400 | 0.670 | 0.219 | 0.068 |  |  |  |  |  |  | 0.277 | 1.319 | 0.207 |
| 362 * 4-day switch * DR | -0.838 | 0.433 | 0.238 | **<0.001** |  |  |  |  |  |  | -0.896 | 0.408 | **<0.001** |
| 441 * 4-day switch * DR | 0.179 | 1.196 | 0.219 | 0.41 |  |  |  |  |  |  | 0.249 | 1.282 | 0.256 |
| 705 * 4-day switch * DR | -0.050 | 0.951 | 0.222 | 0.82 |  |  |  |  |  |  | -0.009 | 0.992 | 0.969 |
| 707 * 4-day switch * DR | -0.910 | 0.403 | 0.232 | **<0.001** |  |  |  |  |  |  | -0.516 | 0.597 | **0.026** |
| 853 * 4-day switch * DR | -0.477 | 0.621 | 0.218 | **0.028** |  |  |  |  |  |  | -0.390 | 0.677 | 0.073 |

**Table S11.**
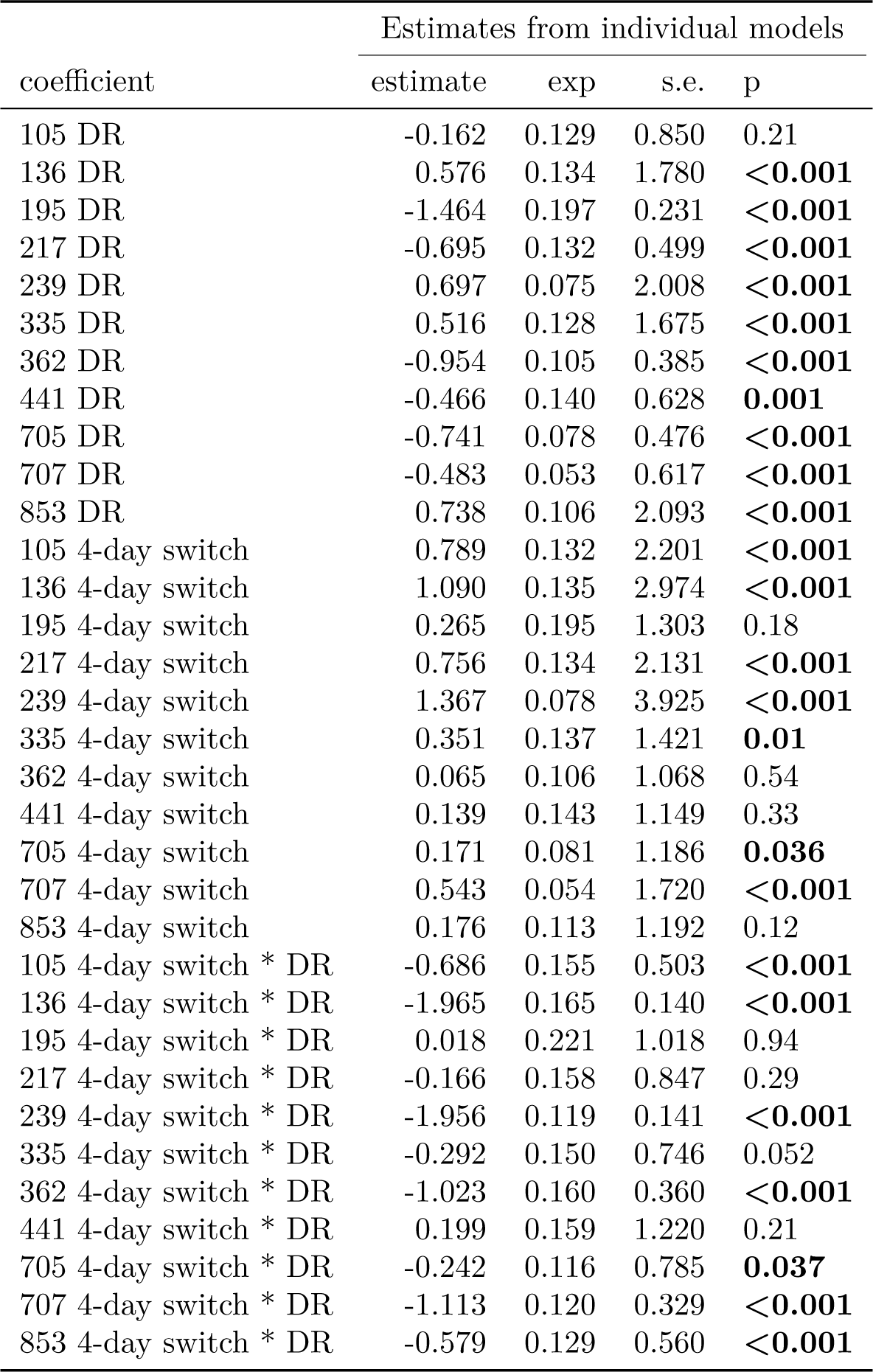
Interval models run within each genotype testing for differential effects of diet in the 4-day switch dietary regime

| coefficient | Estimates from individual models |  |  |  |
| --- | --- | --- | --- | --- |
|  | estimate | exp | s.e. | p |
| 105 DR | -0.162 | 0.129 | 0.850 | 0.21 |
| 136 DR | 0.576 | 0.134 | 1.780 | <**0.001** |
| 195 DR | -1.464 | 0.197 | 0.231 | <**0.001** |
| 217 DR | -0.695 | 0.132 | 0.499 | <**0.001** |
| 239 DR | 0.697 | 0.075 | 2.008 | <**0.001** |
| 335 DR | 0.516 | 0.128 | 1.675 | <**0.001** |
| 362 DR | -0.954 | 0.105 | 0.385 | <**0.001** |
| 441 DR | -0.466 | 0.140 | 0.628 | **0.001** |
| 705 DR | -0.741 | 0.078 | 0.476 | <**0.001** |
| 707 DR | -0.483 | 0.053 | 0.617 | <**0.001** |
| 853 DR | 0.738 | 0.106 | 2.093 | <**0.001** |
| 105 4-day switch | 0.789 | 0.132 | 2.201 | <**0.001** |
| 136 4-day switch | 1.090 | 0.135 | 2.974 | <**0.001** |
| 195 4-day switch | 0.265 | 0.195 | 1.303 | 0.18 |
| 217 4-day switch | 0.756 | 0.134 | 2.131 | <**0.001** |
| 239 4-day switch | 1.367 | 0.078 | 3.925 | <**0.001** |
| 335 4-day switch | 0.351 | 0.137 | 1.421 | **0.01** |
| 362 4-day switch | 0.065 | 0.106 | 1.068 | 0.54 |
| 441 4-day switch | 0.139 | 0.143 | 1.149 | 0.33 |
| 705 4-day switch | 0.171 | 0.081 | 1.186 | **0.036** |
| 707 4-day switch | 0.543 | 0.054 | 1.720 | <**0.001** |
| 853 4-day switch | 0.176 | 0.113 | 1.192 | 0.12 |
| 105 4-day switch * DR | -0.686 | 0.155 | 0.503 | <**0.001** |
| 136 4-day switch * DR | -1.965 | 0.165 | 0.140 | <**0.001** |
| 195 4-day switch * DR | 0.018 | 0.221 | 1.018 | 0.94 |
| 217 4-day switch * DR | -0.166 | 0.158 | 0.847 | 0.29 |
| 239 4-day switch * DR | -1.956 | 0.119 | 0.141 | <**0.001** |
| 335 4-day switch * DR | -0.292 | 0.150 | 0.746 | 0.052 |
| 362 4-day switch * DR | -1.023 | 0.160 | 0.360 | <**0.001** |
| 441 4-day switch * DR | 0.199 | 0.159 | 1.220 | 0.21 |
| 705 4-day switch * DR | -0.242 | 0.116 | 0.785 | **0.037** |
| 707 4-day switch * DR | -1.113 | 0.120 | 0.329 | <**0.001** |
| 853 4-day switch * DR | -0.579 | 0.129 | 0.560 | <**0.001** |

**Table S12.** Models run within each genotype testing for increases in response to a rich diet after a period of DR (long-switch) but within lines that showed starvation only, and with DR as reference category

| coefficient | Estimates from individual models |  |  |  |
| --- | --- | --- | --- | --- |
|  | estimate | exp | s.e. | p |
| 136 rich diet | -0.647 | 0.524 | 0.120 | <**0.001** |
| 239 rich diet | -0.775 | 0.461 | 0.071 | <**0.001** |
| 335 rich diet | -0.570 | 0.566 | 0.115 | <**0.001** |
| 853 rich diet | -0.828 | 0.437 | 0.093 | <**0.001** |
| 136 long switch | 0.113 | 1.120 | 0.132 | 0.39 |
| 239 long switch | 1.483 | 4.406 | 0.103 | <**0.001** |
| 335 long switch | 0.881 | 2.413 | 0.136 | <**0.001** |
| 853 long switch | 1.373 | 3.949 | 0.122 | <**0.001** |

**Table S13.**
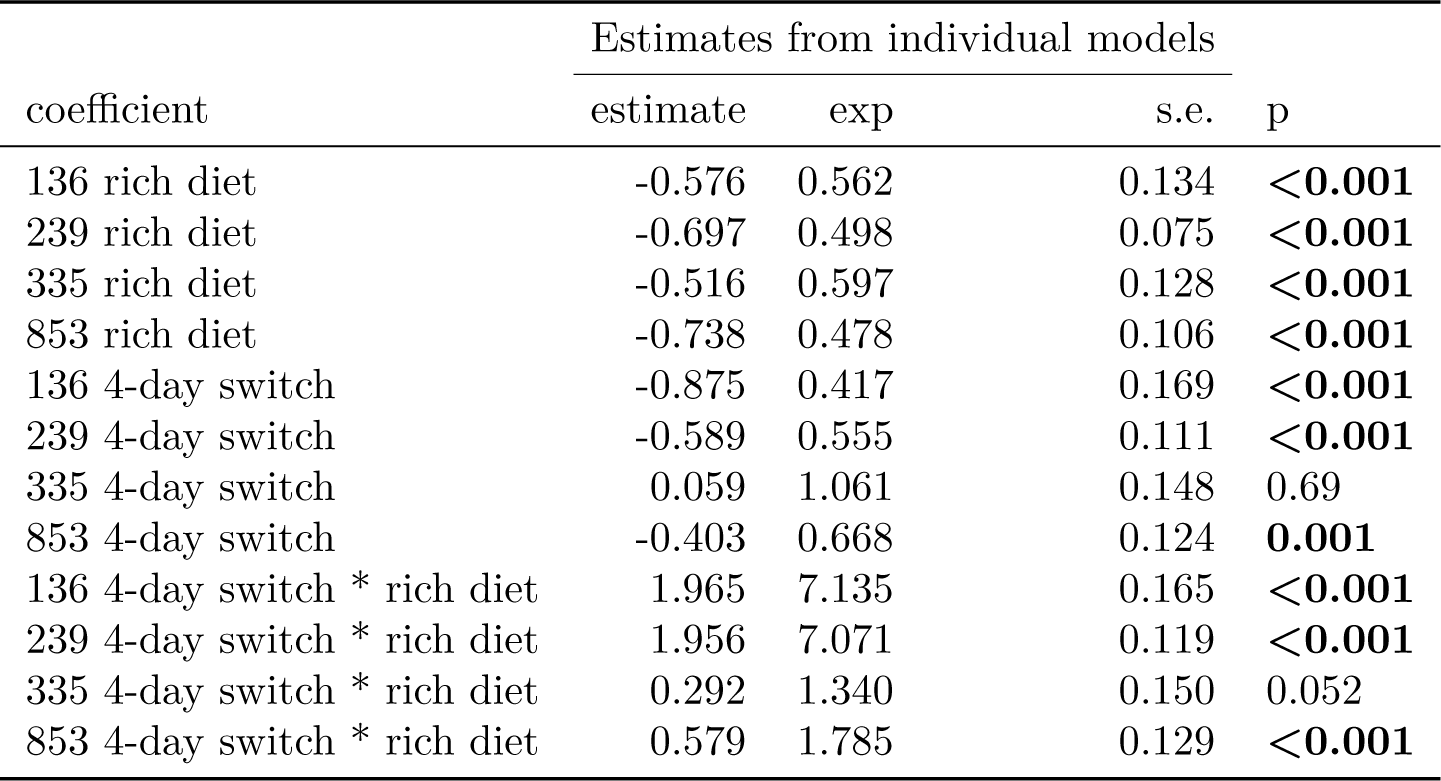
Interval models run within each genotype testing for differential effects of diet in the 4-day switch dietary regime, but within lines that showed starvation only, and with DR as reference category.

| coefficient | Estimates from individual models |  |  |  |
| --- | --- | --- | --- | --- |
|  | estimate | exp | s.e. | p |
| 136 rich diet | -0.576 | 0.562 | 0.134 | **<0.001** |
| 239 rich diet | -0.697 | 0.498 | 0.075 | **<0.001** |
| 335 rich diet | -0.516 | 0.597 | 0.128 | **<0.001** |
| 853 rich diet | -0.738 | 0.478 | 0.106 | **<0.001** |
| 136 4-day switch | -0.875 | 0.417 | 0.169 | **<0.001** |
| 239 4-day switch | -0.589 | 0.555 | 0.111 | **<0.001** |
| 335 4-day switch | 0.059 | 1.061 | 0.148 | 0.69 |
| 853 4-day switch | -0.403 | 0.668 | 0.124 | **0.001** |
| 136 4-day switch * rich diet | 1.965 | 7.135 | 0.165 | **<0.001** |
| 239 4-day switch * rich diet | 1.956 | 7.071 | 0.119 | **<0.001** |
| 335 4-day switch * rich diet | 0.292 | 1.340 | 0.150 | 0.052 |
| 853 4-day switch * rich diet | 0.579 | 1.785 | 0.129 | **<0.001** |

**Table S14.** Linear model of estimates of (log-transformed) fecundity (from Quantifly) in the long-switch dietary treatment. A return to rich conditions from DR, resulted in reduced fecundity, rather than the predicted increase.

| coefficient | Full model |  |  |
| --- | --- | --- | --- |
|  | estimate | s.e. | p |
| rich diet | 2.281 | 0.038 | **<0.001** |
| long switch | -0.367 | 0.037 | **<0.001** |
| 441 | 0.149 | 0.036 | **<0.001** |
| 853 | 0.010 | 0.035 | 0.768 |
| age 45 | 0.040 | 0.041 | 0.339 |
| age 46 | 0.028 | 0.042 | 0.512 |
| age 47 | -0.018 | 0.054 | 0.736 |
| 441 * long switch | -0.195 | 0.050 | **<0.001** |
| 853 * long switch | -0.043 | 0.050 | 0.4 |

**Table S15.** Linear model of estimates of (log-transformed) fecundity (from Quantifly), corrected for number of flies in the cage, in the long-switch dietary treatment. A return to rich conditions from DR, resulted in reduced fecundity, rather than the predicted increase. Note, this correction uses the census after egg-laying and thus overcorrects for mortality. Estimates compared are thus biased upwards, and provide the most sensitive test for an upregulation in response to dietary treatment.

| coefficient | Full model |  |  |
| --- | --- | --- | --- |
|  | estimate | s.e. | p |
| Intercept | 0.374 | 0.048 | **<0.001** |
| long switch | -0.515 | 0.046 | **<0.001** |
| 441 | 0.079 | 0.046 | 0.092 |
| 853 | 0.041 | 0.044 | 0.355 |
| age 45 | 0.049 | 0.052 | 0.349 |
| age 46 | 0.060 | 0.053 | 0.262 |
| age 47 | 0.132 | 0.067 | 0.058 |
| 441 * long switch | -0.158 | 0.063 | **0.017** |
| 853 * long switch | 0.471 | 0.063 | **<0.001** |

**Table S16.** Mixed model (correcting for Cage) of estimates of (log-transformed) fecundity (from Quantifly) in the 4-day switching paradigm. Repeated short-term exposure to DR did not increase, but rather decreased fecundity, relative to a continuous rich diet.

| coefficient | Full model |  |  |
| --- | --- | --- | --- |
|  | estimate | s.e. | p |
| Intercept | 2.924 | 0.057 | <**0.001** |
| DR | -0.642 | 0.030 | <**0.001** |
| 4-day switch | -0.131 | 0.047 | **0.005** |
| 136 | 0.028 | 0.032 | 0.39 |
| 195 | 0.092 | 0.033 | **0.006** |
| 217 | 0.018 | 0.032 | 0.569 |
| 239 | -0.162 | 0.032 | <**0.001** |
| 335 | -0.302 | 0.032 | <**0.001** |
| 362 | -0.092 | 0.033 | **0.005** |
| 705 | -0.570 | 0.032 | <**0.001** |
| 707 | -0.021 | 0.032 | 0.513 |
| 853 | -0.248 | 0.032 | <**0.001** |
| Age 9 | -0.046 | 0.059 | 0.444 |
| Age 10 | -0.073 | 0.054 | 0.171 |
| Age 11 | -0.040 | 0.055 | 0.474 |
| Age 12 | 0.011 | 0.053 | 0.841 |
| Age 13 | -0.086 | 0.054 | 0.11 |
| Age 14 | -0.104 | 0.053 | **0.049** |
| Age 15 | -0.115 | 0.054 | **0.032** |
| Age 16 | -0.139 | 0.053 | **0.009** |
| Age 17 | -0.070 | 0.054 | 0.196 |
| Age 18 | -0.049 | 0.055 | 0.375 |
| Age 19 | -0.202 | 0.056 | <**0.001** |
| Age 20 | -0.212 | 0.056 | <**0.001** |
| Age 21 | -0.367 | 0.072 | <**0.001** |
| 4-day switch * DR | 0.287 | 0.065 | <**0.001** |
| 136 * DR | -0.036 | 0.043 | 0.401 |
| 195 * DR | -0.114 | 0.044 | **0.01** |
| 217 * DR | -0.109 | 0.043 | **0.011** |
| 239 * DR | 0.017 | 0.043 | 0.691 |
| 335 * DR | 0.164 | 0.043 | <**0.001** |
| 362 * DR | 0.072 | 0.043 | 0.09 |
| 705 * DR | 0.422 | 0.043 | <**0.001** |
| 707 * DR | 0.032 | 0.043 | 0.447 |
| 853 * DR | 0.211 | 0.043 | <**0.001** |
| 136 * 4-day switch | -0.014 | 0.066 | 0.829 |
| 195 * 4-day switch | 0.071 | 0.067 | 0.285 |
| 217 * 4-day switch | 0.053 | 0.066 | 0.424 |
| 239 * 4-day switch | -0.029 | 0.066 | 0.656 |
| 335 * 4-day switch | 0.015 | 0.066 | 0.818 |
| 362 * 4-day switch | -0.060 | 0.066 | 0.362 |
| 705 * 4-day switch | 0.219 | 0.066 | **0.001** |
| 707 * 4-day switch | 0.002 | 0.066 | 0.976 |
| 853 * 4-day switch | 0.017 | 0.066 | 0.795 |
| 136 * 4-day switch * DR | 0.215 | 0.092 | **0.02** |
| 195 * 4-day switch * DR | 0.162 | 0.093 | 0.081 |
| 217 * 4-day switch * DR | 0.166 | 0.092 | 0.071 |
| 239 * 4-day switch * DR | -0.048 | 0.092 | 0.601 |
| 335 * 4-day switch * DR | -0.023 | 0.092 | 0.805 |
| 362 * 4-day switch * DR | 0.218 | 0.092 | **0.018** |
| 705 * 4-day switch * DR | -0.422 | 0.092 | <**0.001** |
| 707 * 4-day switch * DR | 0.149 | 0.092 | 0.106 |
| 853 * 4-day switch * DR | 0.112 | 0.092 | 0.225 |

**Table S17.** Mixed model (correcting for Cage) of estimates of (log-transformed) fecundity (from Quantifly), corrected for number of flies in the cage, in the 4-day switching paradigm. Repeated short-term exposure to DR did not increase, but rather decreased fecundity, relative to a continuous rich diet. Note, this correction uses the census after egg-laying and thus overcorrects for mortality. Estimates compared are thus biased upwards, and provide the most sensitive test for an upregulation in response to dietary treatment.

| coefficient | Full model |  |  |
| --- | --- | --- | --- |
|  | estimate | s.e. | p |
| Intercept | 0.812 | 0.056 | **<0.001** |
| DR | -0.640 | 0.030 | **<0.001** |
| 4-day switch | -0.126 | 0.046 | **0.006** |
| 136 | 0.051 | 0.032 | 0.111 |
| 195 | 0.094 | 0.033 | **0.004** |
| 217 | 0.029 | 0.031 | 0.363 |
| 239 | -0.124 | 0.031 | **<0.001** |
| 335 | -0.180 | 0.032 | **<0.001** |
| 362 | -0.057 | 0.032 | 0.08 |
| 705 | -0.556 | 0.031 | **<0.001** |
| 707 | 0.048 | 0.032 | 0.133 |
| 853 | -0.225 | 0.032 | **<0.001** |
| Age 9 | -0.053 | 0.059 | 0.37 |
| Age 10 | -0.089 | 0.053 | 0.094 |
| Age 11 | -0.059 | 0.055 | 0.283 |
| Age 12 | 0.008 | 0.053 | 0.883 |
| Age 13 | -0.083 | 0.053 | 0.12 |
| Age 14 | -0.099 | 0.052 | 0.06 |
| Age 15 | -0.107 | 0.053 | **0.045** |
| Age 16 | -0.120 | 0.053 | **0.022** |
| Age 17 | -0.043 | 0.054 | 0.419 |
| Age 18 | -0.027 | 0.055 | 0.626 |
| Age 19 | -0.161 | 0.055 | **0.003** |
| Age 20 | -0.167 | 0.055 | **0.003** |
| Age 21 | -0.151 | 0.071 | **0.034** |
| 4-day switch * DR | 0.297 | 0.064 | **<0.001** |
| 136 * DR | -0.027 | 0.042 | 0.517 |
| 195 * DR | -0.119 | 0.044 | **0.007** |
| 217 * DR | -0.114 | 0.042 | **0.007** |
| 239 * DR | 0.074 | 0.042 | 0.081 |
| 335 * DR | 0.170 | 0.042 | **<0.001** |
| 362 * DR | 0.064 | 0.042 | 0.129 |
| 705 * DR | 0.418 | 0.042 | **<0.001** |
| 707 * DR | 0.000 | 0.042 | 0.992 |
| 853 * DR | 0.213 | 0.042 | **<0.001** |
| 136 * 4-day switch | 0.003 | 0.065 | 0.969 |
| 195 * 4-day switch | 0.078 | 0.066 | 0.236 |
| 217 * 4-day switch | 0.050 | 0.065 | 0.446 |
| 239 * 4-day switch | 0.076 | 0.065 | 0.243 |
| 335 * 4-day switch | 0.019 | 0.065 | 0.767 |
| 362 * 4-day switch | -0.062 | 0.065 | 0.342 |
| 705 * 4-day switch | 0.217 | 0.065 | **0.001** |
| 707 * 4-day switch | -0.002 | 0.065 | 0.97 |
| 853 * 4-day switch | 0.013 | 0.065 | 0.841 |
| 136 * 4-day switch * DR | 0.178 | 0.091 | 0.051 |
| 195 * 4-day switch * DR | 0.156 | 0.092 | 0.089 |
| 217 * 4-day switch * DR | 0.164 | 0.091 | 0.072 |
| 239 * 4-day switch * DR | -0.060 | 0.091 | 0.511 |
| 335 * 4-day switch * DR | -0.030 | 0.091 | 0.738 |
| 362 * 4-day switch * DR | 0.215 | 0.091 | **0.018** |
| 705 * 4-day switch * DR | -0.431 | 0.091 | **<0.001** |
| 707 * 4-day switch * DR | 0.207 | 0.091 | **0.023** |
| 853 * 4-day switch * DR | 0.098 | 0.091 | 0.283 |

**Table S18.** Effect of returning to a rich diet after a period of DR (long-switch) after ablation of the microbiome (Antibiotics on rich diet is reference)

| coefficient | Full Model |  |  |  |
| --- | --- | --- | --- | --- |
|  | estimate | exp (estimate) | s.e. | p |
| DR on Antibiotics | -0.894 | 0.409 | 0.181 | **<0.001** |
| long switch on Antibiotics | 1.313 | 3.717 | 0.142 | **<0.001** |
| DR | -1.201 | 0.301 | 0.223 | **<0.001** |
| Rich diet | -0.087 | 0.917 | 0.159 | 0.59 |

**Table S19.** Effect of returning to a rich diet after a period of DR (long-switch) with supplementation of water

| coefficient | Full Model |  |  |  |
| --- | --- | --- | --- | --- |
|  | estimate | exp (estimate) | s.e. | p |
| DR with water supplementation | 0.648 | 1.911 | 0.138 | **<0.001** |
| long switch with water supplementation | 0.385 | 1.470 | 0.126 | **0.002** |

**Table S20.** Effect of returning to a rich diet after a period of DR (long-switch) with flies in isolation in vials. Test from chi-square tests on proportions by age.

| Age | Dead |  |  |  | Mortality rate |  |  |  | P value from proportion test |  |
| --- | --- | --- | --- | --- | --- | --- | --- | --- | --- | --- |
|  | DR | Rich to DR | Rich | DR to Rich (long switch) | DR | Rich to DR | Rich | DR to Rich (long switch) | Rich versus DR | Rich versus long switch |
| 28 | 0 | 0 | 5 | 16 | 0.000 | 0 | 0.100 | 0.320 | 0.063 | **0.014** |
| 30 | 0 | 0 | 8 | 5 | 0.000 | 0 | 0.178 | 0.152 | **0.006** | 1 |
| 32 | 1 | 0 | 16 | 13 | 0.020 | 0 | 0.432 | 0.464 | **<0.001** | 0.997 |
| 34 | 1 | 0 | 8 | 4 | 0.020 | 0 | 0.381 | 0.267 | **<0.001** | 0.72 |
| 36 | 1 | 0 | 8 | 2 | 0.020 | 0 | 0.667 | 0.200 | **<0.001** | 0.079 |
| 38 | 4 | 0 | 2 | 2 | 0.087 | 0 | 0.500 | 0.250 | 0.102 | 0.829 |
| 40 | 3 | 0 | 0 | 3 | 0.071 | 0 | 0.000 | 0.500 | 1 | 1 |
| 42 | 2 | 0 | 0 | 3 | 0.051 | 0 | 0.000 | 1.000 | 1 | 0.505 |

**Table S21.** Effect of switching from DR to rich food every four days (4-day switch) in males

| coefficient | Full Model |  |  |  |
| --- | --- | --- | --- | --- |
|  | estimate | exp (estimate) | s.e. | p |
| DR | 0.362 | 1.437 | 0.117 | **0.002** |
| 4-day switch | 0.410 | 1.507 | 0.120 | **0.001** |
| 4-day switch * DR | -0.768 | 0.464 | 0.129 | **<0.001** |

